# STRIPAK complex defects result in pseudosexual reproduction in *Cryptococcus neoformans*

**DOI:** 10.1101/2025.04.08.647827

**Authors:** Patricia P. Peterson, Sarah Croog, Yeseul Choi, Sheng Sun, Joseph Heitman

**Author notes:** Corresponding author: (JH).

## Abstract

STRIPAK is an evolutionarily conserved signaling complex that coordinates diverse cellular processes across fungi and animals. In the human fungal pathogen *Cryptococcus neoformans*, STRIPAK was recently shown to play critical roles in maintaining genome stability and controlling both sexual and asexual development. In *Cryptococcus*, sexual reproduction is closely linked to virulence, and our findings demonstrate that the STRIPAK complex plays key roles in both processes. Here, we further investigate the specific roles of the STRIPAK catalytic subunit Pph22 and its regulatory partner Far8 during sexual development. We show that while *pph22*Δ mutants are defective in α-**a** sexual reproduction, exhibiting impaired meiotic progression and a failure to produce viable spores, the deletion of *PPH22* resulted in exclusive pseudosexual reproduction, with progeny inheriting nuclear genomes solely from the wild-type parent. Overexpression of *PPG1*, a related phosphatase, rescued growth and developmental defects in *pph22*Δ mutants, and restored the preference for α-**a** sexual reproduction over pseudosexual reproduction during mating, suggesting functional redundancy within the STRIPAK signaling network. Furthermore, deletion of *FAR8*, another component of the STRIPAK complex, also led to a high rate of pseudosexual reproduction during α-**a** sexual mating, reinforcing the role of STRIPAK in modulating reproductive modes in *C. neoformans*, possibly through regulating nuclear inheritance and meiotic progression. Together, these findings highlight the distinct contributions of STRIPAK to sexual reproduction in *C. neoformans* and suggest that disruptions of this complex affect genome integrity and inheritance mechanisms, with broader implications for fungal adaptation and pathogenesis.

## Introduction

In eukaryotic cells, cellular processes are orchestrated by the coordinated activities of protein kinases and phosphatases. Among these, protein phosphatase 2A (PP2A) is a pivotal serine/threonine phosphatase that governs essential functions in cell growth, metabolism, stress responses, proliferation, and differentiation [1–5]. PP2A operates as a heterotrimeric complex, consisting of a scaffold A subunit, a catalytic C subunit, and a regulatory B subunit, which dictates substrate specificity and pathway regulation [6–8]. One key regulatory B subunit is striatin, a core component of the striatin-interacting phosphatase and kinase (STRIPAK) complex, a highly conserved signaling complex that integrates phosphatase and kinase activity to govern developmental and cellular processes across eukaryotes [1, 9–11]. STRIPAK assembly begins with the formation of the PP2A-striatin holoenzyme, which serves as a platform for recruiting additional components to build a larger signaling module [1, 12]. STRIPAK complexes have been identified and characterized in yeasts, filamentous fungi, and mammals, including humans, underscoring their evolutionary conservation. While STRIPAK is broadly conserved across eukaryotes, its functional roles can be context dependent, varying across different organisms and cellular environments [11, 13].

In fungi, STRIPAK is required for both sexual and asexual development, influencing key processes such as cell cycle progression, conidiation, cell fusion, and sporulation [10, 14–17]. In yeasts, STRIPAK-like complexes containing PP2A components are important in the pheromone response pathway and cytokinesis during the sexual cycle [18–20]. In filamentous ascomycetes, STRIPAK homologs have been extensively studied for their roles in sexual and asexual propagation [15, 17, 21–23]. In these species, STRIPAK is essential for the formation of mature fruiting bodies, which are highly complex multicellular structures that arise during sexual reproduction, as well as for asexual growth through conidiation [24, 25]. Similarly, in plant pathogenic fungi such as *Magnaporthe oryzae, Collectrichum graminicola,* and *Fusarium* species, STRIPAK is linked to developmental transitions that enhance virulence, affecting the ability of these pathogens to infect host tissues [26–29]. More recently, STRIPAK has been characterized in the basidiomycete pathogens *Ustilago maydis* and *Cryptococcus neoformans,* where it plays conserved roles in vegetative growth and sexual reproduction [11, 30]. In *Cryptococcus*, STRIPAK also contributes to genome stability, virulence, and pathogenesis, highlighting its broader significance beyond development.

*C. neoformans* is a human fungal pathogen in which sexual reproduction has been linked to virulence. Among natural isolates, the α mating type is more prevalent and has been associated with increased virulence, while sexual basidiospores and desiccated yeast cells serve as infectious propagules [31–34]. The dimorphic transition of budding yeast to hyphal growth after mating has also been implicated in pathogenesis, and the formation of sexual structures in environmental niches may enhance fungal survival [35]. Mating between cells of the two opposite mating types, α and **a**, begins with cell-cell fusion, followed by hyphal filamentation, and the formation of a basidium, where nuclear fusion and meiosis occur. After meiosis, nuclei are partitioned into budding spores produced from the basidium, which undergo mitotic division to produce spore chains [36]. Notably, each basidium represents a single meiotic event, and the resulting basidiospores, which form in chains, exhibit distinct adjacent genotypes due to the segregation of alleles during meiosis [37]. This enables the generation of diverse offspring from a single mating event, which may enhance dispersal and dissemination in the environment. Altogether, this sexual cycle contributes not only to genetic diversity but also to environmental persistence and ecological adaptability.

In addition to conventional α-**a** sexual reproduction, *Cryptococcus* can undergo pseudosexual reproduction, an alternative reproductive strategy that results in uniparental nuclear inheritance following mating [38]. Unlike classic meiosis, where both parental nuclei contribute genetic material via recombination to produce haploid progeny, pseudosexual reproduction involves the preferential retention of one parental nucleus while the other is lost or excluded, leading to meiotic progeny whose nuclear genome is genetically identical to one parent and thus lack meiotic recombination. Mitochondrial uniparental inheritance (UPI) also occurs during this process, with mitochondria preferentially inherited from the *MAT***a** parent [39]. This process shares striking similarities with hybridogenesis, a reproductive mode observed in some animals where hybrid offspring selectively eliminate one parental genome during gametogenesis, ensuring uniparental inheritance in subsequent generations [40]. In *Cryptococcus*, pseudosexual reproduction may facilitate progeny survival and expansion in environments where compatible mating partners are scarce, particularly when high genetic divergence or karyotype variation limits traditional mating. Despite its potential role in adaptation and persistence, the molecular mechanisms underlying this process, and the genetic factors involved, remain largely unknown.

In this study, we investigated the roles of Pph22 and Far8, two key components of the STRIPAK complex, which are critical for *C. neoformans* sexual development. Genotyping and whole-genome sequencing of progeny from *pph22*Δ × wild-type crosses revealed that reproduction occurs exclusively through pseudosexual reproduction, with uniparental nuclear inheritance derived solely from the wild-type parent. Fluorescence microscopy further demonstrated that while *pph22*Δ nuclei can migrate into the basidium and initiate meiosis, they fail to generate viable spores, implicating *PPH22* in meiotic progression. Additionally, overexpression of *PPG1*, a functionally related phosphatase, suppressed defects in vegetative growth, stress responses, and sexual growth caused by loss of *PPH22*, highlighting potential compensatory mechanisms within the STRIPAK network. Notably, we found that pseudosexual reproduction also occurs at a high rate in *far8*Δ mutant x wild type crosses, further supporting a role for STRIPAK in coordinating nuclear inheritance and meiotic processes. These findings establish STRIPAK as a key regulatory network in *Cryptococcus* reproduction and suggest that disruptions to this complex may broadly influence genetic stability and inheritance mechanisms in this fungal pathogen.

## Results

### *pph22*Δ mutants exhibit reduced mating efficiency and abnormal sexual structures

We previously showed that deletion of *PPH22* causes severe mating defects during α-**a** sexual reproduction, as well as in a self-fertile α/**a** diploid strain, leading to significantly delayed sexual development [30]. To further investigate the dynamics of *pph22*Δ mutants during mating, *pph22*Δ strains were crossed with a wild-type partner of the opposite mating type and incubated on MS medium. To overcome the growth defects of *pph22*Δ on nutrient-limited MS media, mating was facilitated by pre-growing mutant cells before adding a smaller amount of wild-type cells, as described previously [30]. After six weeks of incubation, wild-type crosses displayed extensive hyphal growth, basidia formation, and basidiospore production. In contrast, *pph22*Δ x wild type mating patches showed only limited signs of sexual development (Fig 1A). Most patches lacked visible sexual structures, though occasional hyphae and basidia were observed. Basidia were often bald or produced few spores, and many spores appeared collapsed rather than forming the elongated spore chains characteristic of wild-type mating.

**Figure 1.**
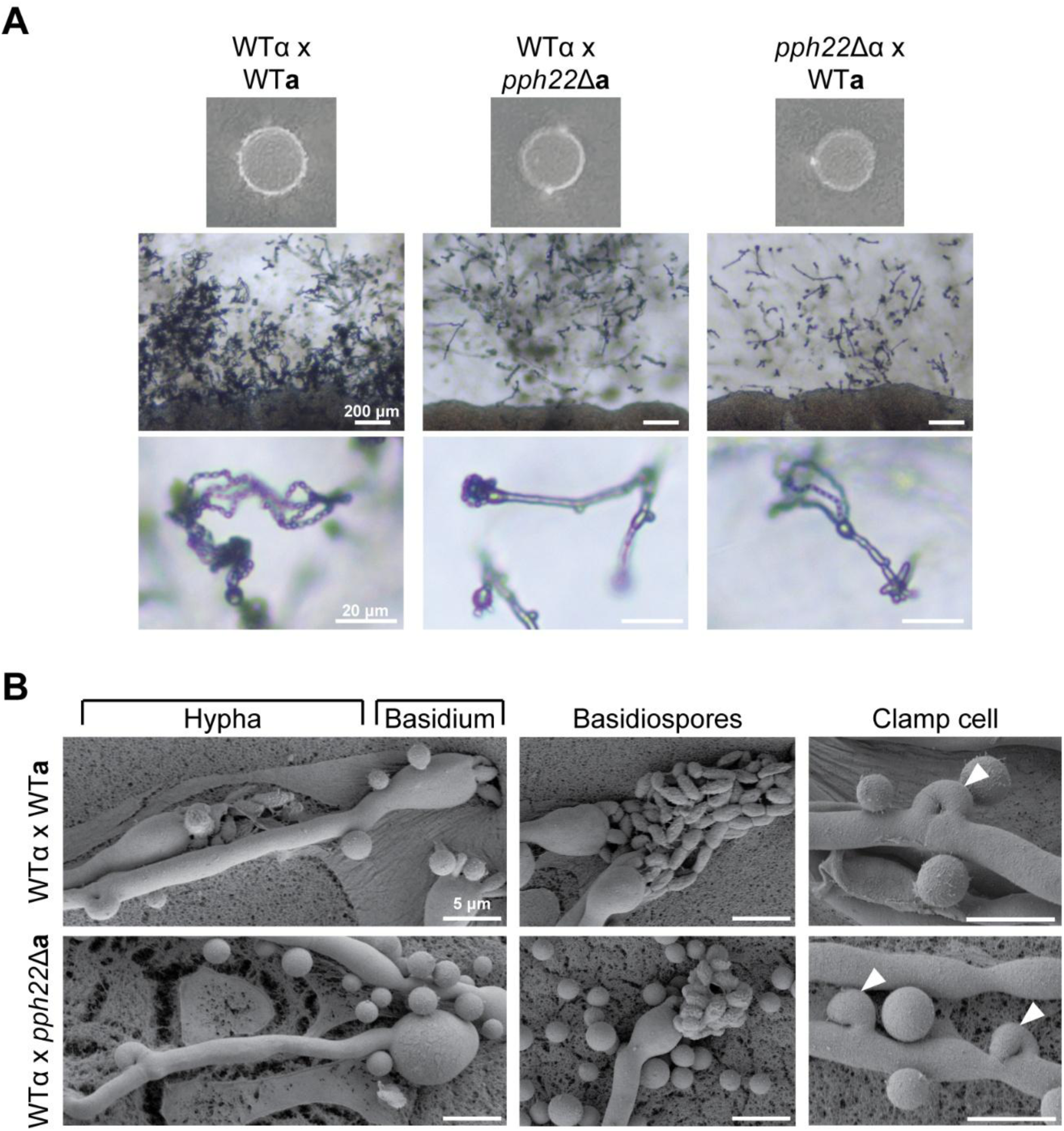
*pph22*Δ x wild type crosses exhibit unusual sexual development. A) Mating efficiency of *pph22*Δ strains crossed with wild-type of the opposite mating type (H99α or KN99**a**). Images were taken between 4 to 6 weeks of incubation on MS plates. Scale bars in the middle and bottom panels are 200 μm and 20 μm, respectively. B) Scanning electron microscopy (SEM) of sexual structures in WT x WT and *pph22*Δ x WT crosses showing examples of a hyphal filament terminating in a basidial head and basidiospores emerging from a basidium. The clamp cells (white arrowheads) in the wild-type cross are fused to the subapical cell in the elongating hypha. In *pph22*Δ x WT crosses, both fused and unfused clamp cells were often observed along the same hypha. Scale bars in each panel represent 5 μm.

To observe morphological differences in sexual structures between *pph22*Δ and wild-type crosses at a higher resolution, scanning electron microscopy (SEM) of the mated strains was performed. In *pph22*Δ x wild type crosses, many basidia terminating from the end of a hyphal branch were considerably rounded and enlarged compared to the wild-type cross and did not produce spores (Fig 1B). When spores were observed, they often had a collapsed appearance, similar to what was seen by light microscopy (Fig 1A). During α-**a** sexual reproduction in the wild type x wild type cross, the clamp cells, which are critical for faithful segregation of nuclei in the dikaryotic hyphae [41, 42], were fused to the subapical cell in the elongating hyphal branch. In mated *pph22*Δ x wild type, both fused and unfused clamp cells could be observed along the same hypha, suggesting a potential defect in nuclear distribution or clamp cell fusion. Due to the observed defects in mating, we quantified cell–cell fusion frequency by incubating crosses between *pph22*Δ*::NAT* and wild type::*NEO* strains on MS medium for three days. After incubation, cells were harvested and plated on YPD supplemented with either NAT, NEO, or both NAT and NEO. Crosses between wild type::*NAT* and wild type::*NEO* strains were included as a control. Fusion events were quantified by counting colony-forming units (CFUs) on YPD+NAT+NEO plates, which represent dual-resistant fusion products. Fusion frequency was calculated as the number of CFUs on YPD+NAT+NEO divided by the sum of CFUs on YPD+NAT and YPD+NEO, representing the total recovered population. A significant reduction in fusion frequency was observed in *pph22*Δ**a** × WTα and *pph22*Δα × WT**a** crosses compared to the wild-type control (Fig S1A). There was a significant reduction in cell-cell fusion frequency in *pph22*Δ**a** x WTα and *pph22*Δα x WT**a** crosses (5.1% to 5.7% of WT x WT). Taken together, these results suggest that *PPH22* is critical for α-**a** sexual mating and proper sexual development.

### Progeny from *pph22*Δ x wild type crosses are produced exclusively through pseudosexual reproduction

To further investigate the reproductive mode of *pph22*Δ x WT crosses, spores were dissected from individual basidia for phenotypic and genotypic analysis. *pph22*Δ mutant strains have severe growth defects and form significantly smaller colonies compared to wild type. Germinated spores from *pph22*Δ x WT formed colonies that were fast growing and uniform in size on YPD (Fig S1B). Streaking onto YPD and YPD+NAT (selecting for the *pph22*Δ*::NAT* mutation) showed that all dissected spores were NAT-sensitive, indicating the absence of *pph22*Δ mutants among the recovered progeny, potentially suggesting that the spores originated from unisexual reproduction of the wild-type parental strain [43]. To investigate this possibility, spores were dissected from basidia of two independent *pph22*Δ**a** × WTα crosses and one *pph22*Δα × WT**a** cross to yield viable progeny. PCR genotyping of the *STE20* gene in the mating-type locus *(MAT*) revealed that all progeny from each individual basidium contained either the *MAT***a** or the *MAT*α allele, which strictly corresponded to the allele from the wild-type parent (Table 1, Fig S1C). PCR of the *COX1* gene also indicated that the mitochondria was predominantly inherited from the *MAT***a** parent, consistent with mitochondrial uniparental inheritance (mito-UPI) in *Cryptococcus* [39, 44]. *MAT*α mitochondrial inheritance was observed in *pph22*Δ**a** × WTα crosses at a frequency of 32% (7 out of 22 basidia), which is higher than the expected frequency of *MAT*α mitochondria leakage of around 5% to 10% [45]. However, six of these basidia originated from the same mating spot, leaving open the possibility that the inherited *MAT*α mitochondria in the progeny resulted from a single mating event. Similarly, in random spore dissection, 15 out of 65 germinated progeny possessed *MAT*α mitochondria, but were dissected from a single mating spot. It is unclear whether the *pph22*Δ mutation leads to increased *MAT*α mitochondrial inheritance. These results suggest that progeny from *pph22*Δ × WT inherited only the wild-type parental nuclei. Additionally, the presence of mitochondria from the *MAT***a** parent in *MAT*α progeny indicates that these progeny are not the result of unisexual reproduction. These observations demonstrate that the mode of reproduction in *pph22*Δ × WT crosses is pseudosexual, during which cells of the opposite mating type fuse but the nuclear content of only one parental nucleus is transmitted to the progeny [38].

**Table 1.**
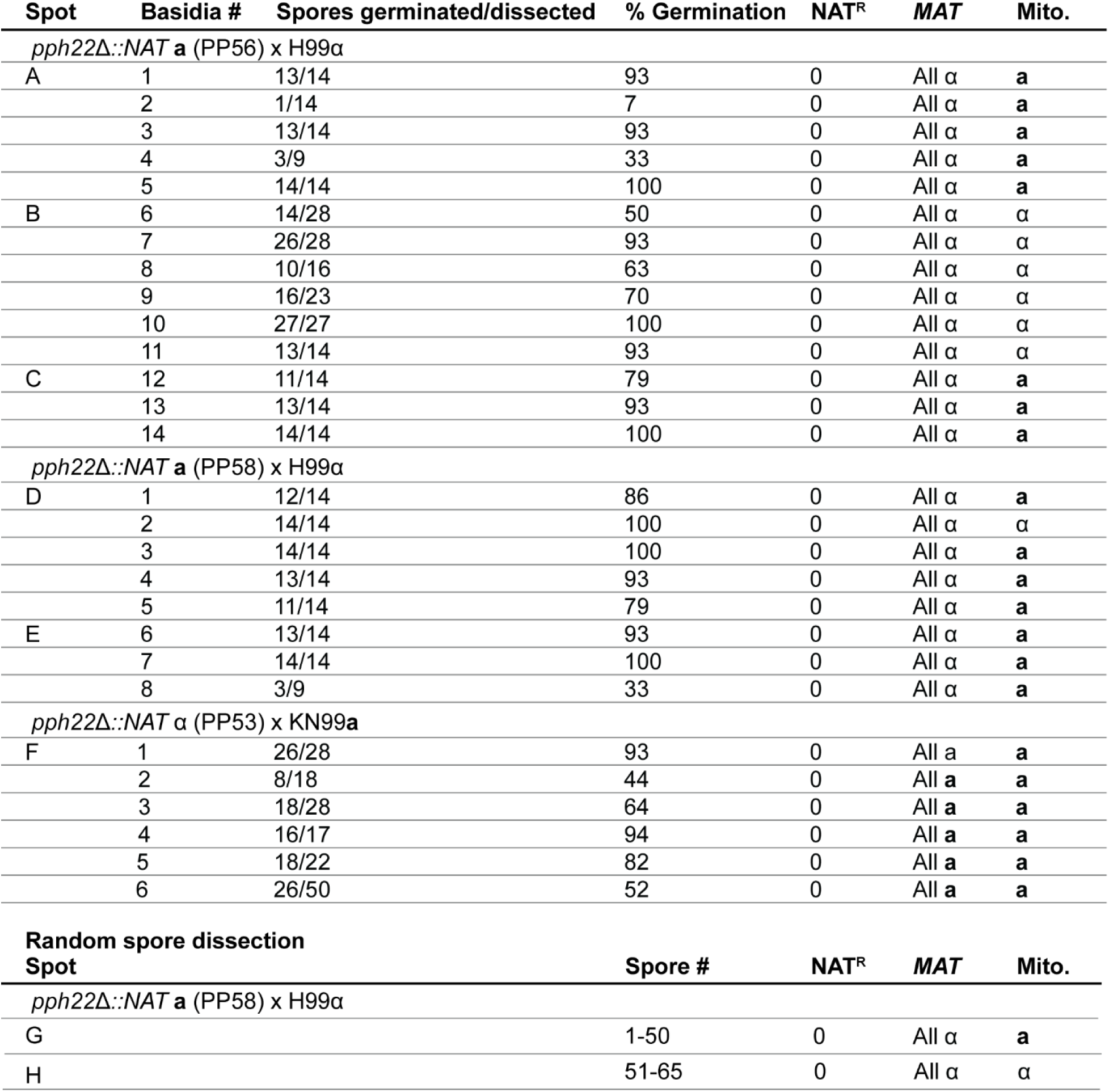
Genotyping analysis of spores from individual basidia and from random spore dissection in *pph22*Δ x wild type crosses. Spot refers to the individual mating spot from which those basidia were dissected in the indicated crosses, with each spot being an independent experiment. Mito. refers to mitochondria.

Following PCR genotyping assays, whole-genome sequence analysis was conducted to determine if there was any evidence of meiotic recombination in the F1 progeny from *pph22*Δ**a** × WTα. The *pph22*Δ strains utilized in this study were obtained from dissection of a KN99**a/**α *PPH22/pph22*Δ heterozygous diploid mutant and are congenic with H99 [30, 46]. Besides the mating-type locus on chromosome 5, there are two other genomic regions on chromosomes 11 and 14 that differ between H99α and KN99**a** background strains [47, 48]. Twelve progeny from a single basidia (Table 1, spot D basidium 1) were subjected to Illumina whole-genome sequencing and reads were aligned to the H99α reference genome for variant calling and SNP analysis (Fig 2). The KN99**a** *pph22*Δ parental strain and the KN99**a/**α diploid strain (CnLC6693) from which it was derived served as controls. Variant analysis confirmed that only the H99α nuclear genome was inherited in the progeny, as evidenced by the absence of SNPs at the indicated polymorphic loci. Alignment of reads from the KN99 parental strains and progeny to the H99α mitochondrial genome also confirmed the presence of the intron in the KN99**a** *COX1* gene, supporting the results from PCR genotyping. Collectively, the lack of meiotic recombination supports that the progeny from mating of *pph22*Δ × WT arose through a pseudosexual reproductive process and not via α-**a** sexual reproduction.

**Figure 2.**
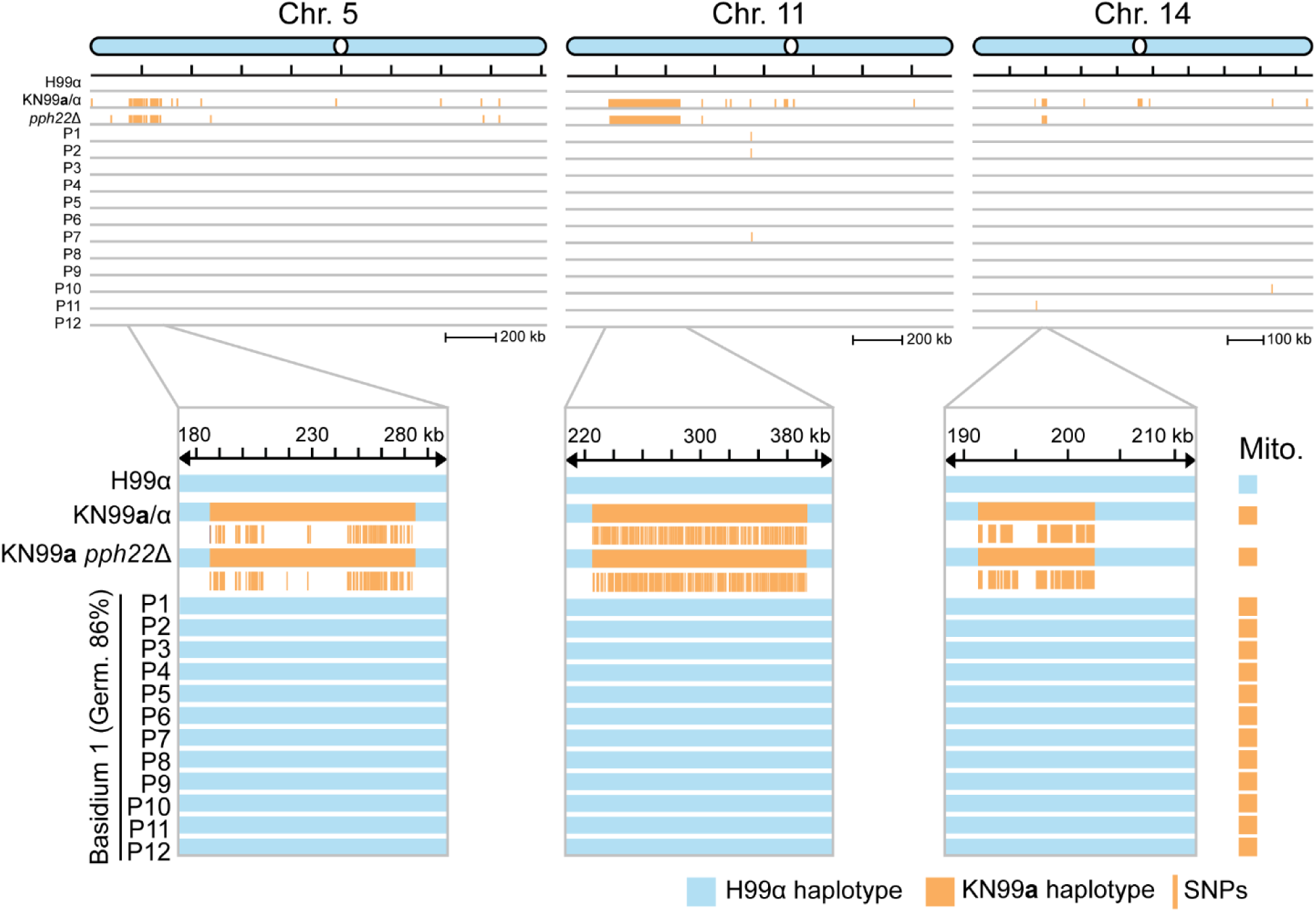
Progeny from *pph22*Δ x wild type exhibit uniparental nuclear inheritance from H99α and lack evidence of meiotic recombination. Whole-genome sequencing followed by SNP analysis revealed no contribution of the KN99**a** *pph22*Δ parental genome, as evidenced by the absence of SNPs compared to the H99α reference genome. The three known polymorphic regions distinguishing H99 and KN99 background strains are located on chromosomes 5 (the mating-type locus), 11, and 14. These regions are highlighted in the lower panels, with the chromosomal coordinates labeled accordingly. The KN99**a**/KN99α diploid strain (CnLC6683), from which KN99**a** *pph22*Δ was obtained, was used as a control for variant calling. The few SNPs identified in the progeny were restricted to nucleotide repeat regions. Mitochondrial haplotypes were determined based on the presence or absence of an intron in the *COX1* gene from whole-genome sequencing reads. “Germ.” stands for the germination frequency of the indicated basidium and “P” stands for progeny.

### Uniparental nuclear inheritance observed after mating via fluorescence microscopy

Previous studies with fluorescently marked nuclei provided evidence that pseudosexual reproduction in *Cryptococcus* results from loss of one parental nucleus during hyphal branching following mating [38]. A similar approach was taken to investigate nuclear migration during mating in *pph22*Δ × WT crosses. The gene encoding the nucleolar protein Nop1 was fused with GFP or RFP (mCherry) and introduced ectopically into the safe haven region of *pph22*Δ**a** (*NOP1-GFP-NEO)*, KN99**a** (*NOP1-GFP-NEO)*, and H99α (*NOP1-mCherry-HYG*) [49].

Homologous recombination at the target locus was confirmed by PCR genotyping. *pph22*Δ and KN99**a** strains expressing Nop1-GFP were crossed with H99α expressing Nop1-RFP on MS media and monitored for signs of mating (Fig S2A). In wild-type crosses, live-cell imaging revealed hyphae containing both GFP- and RFP-tagged proteins following cell-cell fusion, maintaining a stable dikaryotic state throughout hyphal elongation and branching, with green and red fluorescent signals overlapping or appearing in close proximity (Fig S2B). In the basidial heads, the final stages of the sexual cycle were observed, including karyogamy, where the two parental nuclei fused, followed by meiosis and subsequent inheritance of fluorescent tags from both parents in the haploid basidiospores. In total, 27 basidia from two independent crosses were observed to produce spores that contained both fluorescent markers.

Nuclear dynamics in *pph22*Δ crosses were notably distinct from those in wild-type crosses. As expected, *pph22*Δ**a** x H99α exhibited decreased mating efficiency, but spore production was still observed after six to eight weeks of incubation (Fig S2A). Fluorescence imaging showed that both red and green nuclear markers initially coexisted within the same hyphal compartment and could be tracked along the elongating hyphal branch. However, at certain branch points, the green *pph22*Δ nucleus was selectively lost, allowing only the red wild-type nucleus to undergo meiosis in the terminating basidium (Fig S2C). Among 40 basidia analyzed from four independent crosses, 15 did not produce spores. Of these, 12 displayed both red and green signals in the basidium head, while the remaining three showed only the red signal. The other 25 basidia successfully produced spores, with 20 containing only the signal from the wild-type parent. This finding aligns with previous work suggesting that hyphal branching may facilitate nuclear exclusion, which can be followed by endoreplication and meiosis of the retained nucleus [38].

In contrast, there were also cases in crosses of *pph22*Δ x WT where the wild-type nucleus was lost following hyphal branching, after which the resulting sexual structures exhibited abnormal morphology (Fig 3). Several observations suggested that, after the loss of the red wild-type signal in a hyphal branch, the green *pph22Δ* nucleus could undergo endoreplication followed by nuclear division, resulting in two green nuclei within a single hyphal compartment, indicating a mechanism by which a dikaryotic-like state could be reestablished (Figure 3A). There were also instances where, following wild-type nuclear exclusion, only the *pph22*Δ fluorescent signal was inherited in the basidiospores, which appeared collapsed onto the basidium head (Fig 3B, 3C). Notably, five basidia that produced spores exhibited only the green signal in the progeny., suggesting that the *pph22*Δ mutant nucleus is capable of migrating into the basidium, undergoing diploidization, and potentially entering the meiotic process.

**Figure 3.**
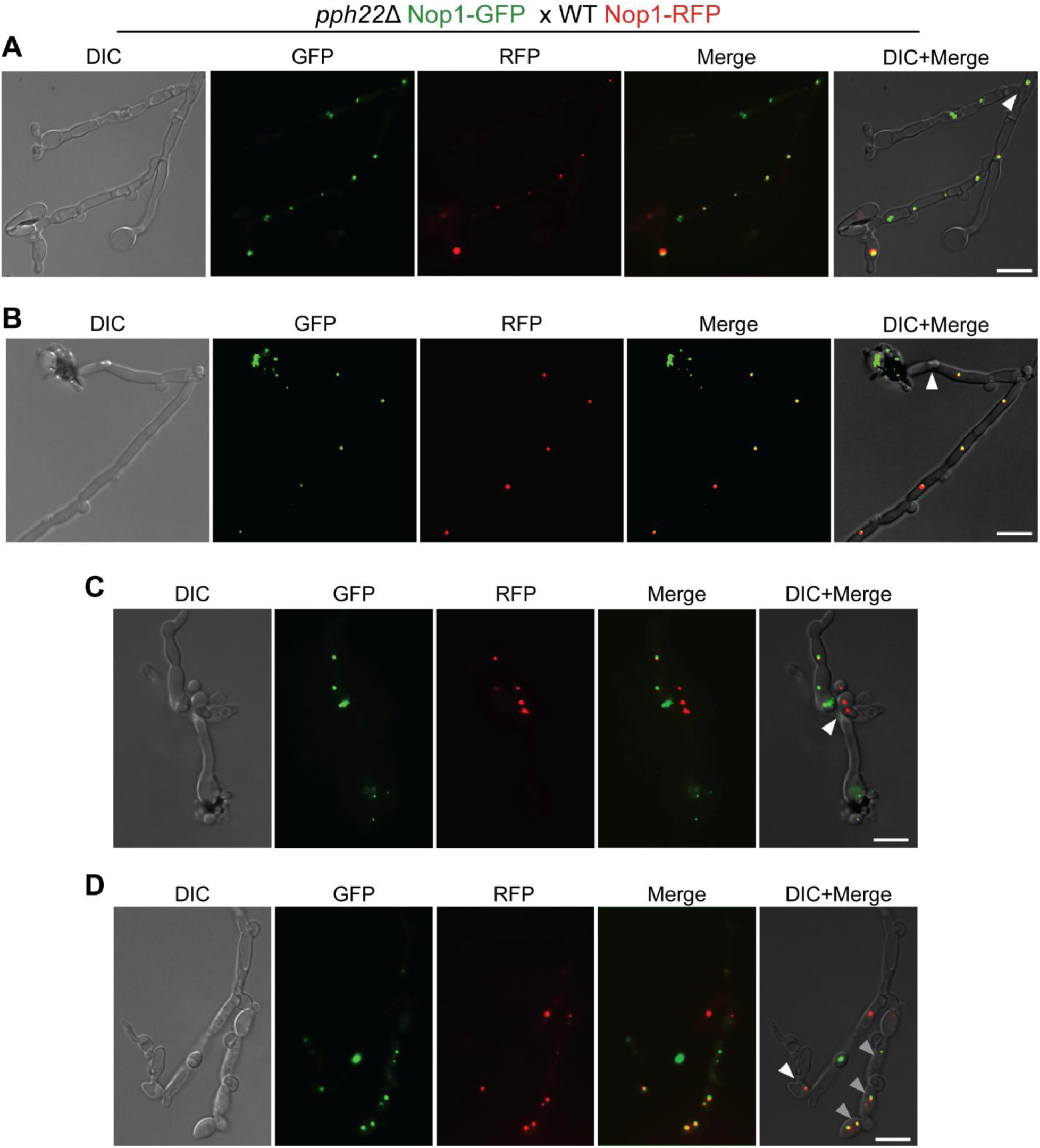
Fluorescence microscopy depicts loss of one parental nucleus following hyphal branching during pseudosexual reproduction. A) Loss of the wild-type nucleus at a hyphal branch point in *pph22*Δ Nop1-GFP x WT Nop1-RFP. B) and C) Inheritance of only the *pph22*Δ nucleus in basidiospores, which appear collapsed on the basidial head and have an irregular morphology, is observed after loss of the WT nucleus in a hyphal compartment (B) and following branching (C). D) Loss of the wild-type nucleus and retention of the *pph22*Δ nucleus results in severely abnormal hyphal development (white arrowhead). In another hyphal branch, both nuclei have undergone division in the dikaryon without segregating into separate hyphal cells (grey arrowheads). The white arrowheads indicate the point in the hypha where one parental nucleus is lost. The scale bar in each panel represents 5 μm.

However, despite this apparent progression, the consistent inability to recover *pph22*Δ mutant progeny from any dissection of *pph22*Δ x WT crosses, indicates a meiotic or post-meiotic defect that prevents successful germination. There was also evidence that maintenance of only the *pph22*Δ nucleus negatively impacted hyphal morphology, significantly altering both the shape and structure (Fig 3D). Additionally, in another hyphal branch, there were two nuclei detected for each color to yield four nuclei in individual cells, suggesting a failure in nuclear sorting or segregation following DNA replication. Together with spore genotyping analyses, these findings suggest that: 1) pseudosexual reproduction may occur through the loss of one parental nucleus during hyphal branching and 2) the *pph22*Δ mutation disrupts normal sexual development, including nuclear migration, hyphal morphology, and sporulation.

### α-a sexual reproduction can be restored in *pph22*Δ *suppressor* mutants

*pph22*Δ strains can accumulate spontaneous suppressor mutations that restore mating, hyphal growth, basidia production, and sporulation nearly to the wild-type level (Fig 4A) [30]. To determine if the *pph22*Δ *suppressor* (*sup)* mutant strains could also suppress pseudosexual reproduction, F1 progeny from *pph22*Δ *sup* x WT crosses were analyzed. In one such cross, germinated spores containing the *NAT* drug-resistance marker were recovered during dissection, in contrast to strictly NAT-sensitive *pph22*Δ x WT progeny, providing evidence the suppressor mutation could restore *pph22*Δ spore viability following α-**a** sexual mating, although the resulting colonies were notably smaller in size (Fig 4B). The frequency of *pph22*Δ::*NAT* among the progeny was 45%, consistent with the expected segregation of a single selectable allele. However, in crosses with two other independently derived *pph22*Δ *sup* strains, no NAT resistant progeny were recovered, suggesting incomplete suppression of *pph22*Δ mutant defects following mating in those strains.

**Figure 4.**
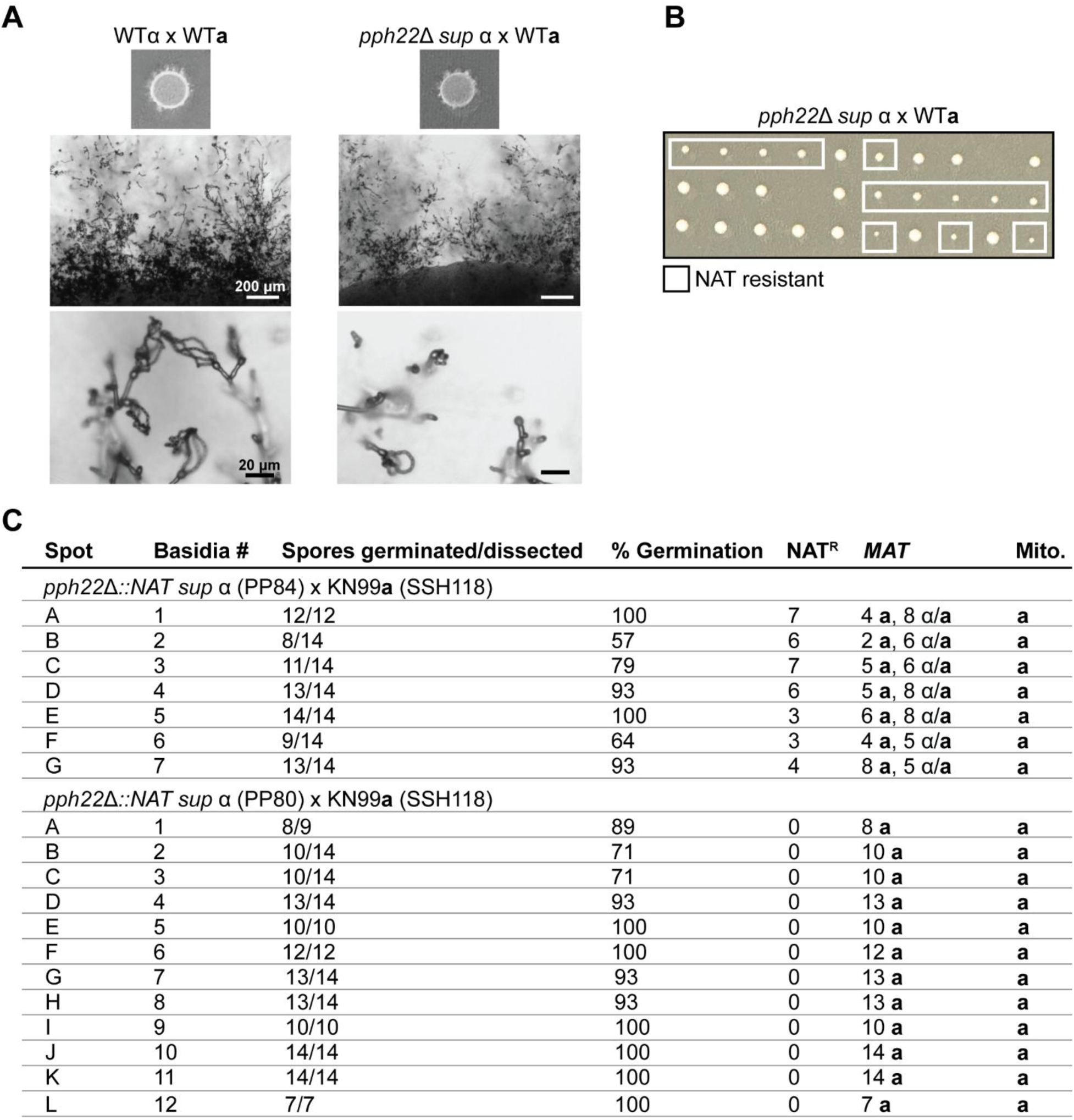
The *pph22*Δ *suppressor* strains can restore α-**a** sexual mating in crosses with wild type. A) Mating efficiencies in wild-type and *pph22*Δ *sup* crosses. Hyphal production and elongation, and sporulation were observed to similar extents. B) Germination of spores dissected from *pph22*Δ *sup* x WT after two days of incubation on YPD. Each row depicts spores from separate basidia. Colonies that grew on YPD+NAT (containing the *pph22*Δ*::NAT* mutant allele) are indicated by the white boxes. C) Genotyping analysis of progeny dissected from *pph22*Δ *sup* strains crossed with the KN99**a** strain bearing recombinant mitochondria (SSH118). Each spot is from a separate mating patch and is considered an independent experiment.

We next investigated mating type and mitochondrial genome segregation in *pph22*Δ *sup* x WT crosses. All *pph22*Δ *sup* isolates that were obtained in our previous study possess the *MAT*α allele and therefore were crossed with KN99**a**. This led to a complication in using the same PCR genotyping strategy to determine how the mitochondria was inherited in the progeny from these crosses, as KN99**a** and KN99α background strains have identical mitochondria genomes. Therefore, *pph22*Δ *sup* strains were crossed with the KN99**a** strain SSH118, which carries a recombinant mitochondrial genome, and restriction fragment length polymorphism (RFLP) analysis was used to identify the parental source of the mitochondrial genome in the progeny (Fig 4C, S3). In each *pph22*Δ *sup* x WT cross, each basidium that was dissected came from different mating spots, so that the progeny analyzed were the result of independent mating events. The *MAT* allele was determined through PCR genotyping of the *STE20* gene, as in previous experiments, and mitochondrial type was identified through PCR amplification of *COX1* followed by BsrI digestion and RFLP analysis (Fig S3B, S3C, S3D). In two independent crosses, both **a** and α mating types were found among the genotyped basidia and, interestingly, many progeny were heterozygous for the *MAT* allele (Fig 4C, S3E). However, in a third independent cross with a different *pph22*Δ *sup* strain, only progeny of one mating type were recovered, consistent with pseudosexual reproduction. The fact that progeny exhibiting *MAT* heterozygosity also came from basidia with high germination rates suggests that this cannot be explained by aneuploidy alone, as an imbalance in chromosome number in aneuploid spores would lead to reduced germination rates.

Flow cytometry analysis of progeny heterozygous for the *MAT* locus revealed that increased DNA content correlated with the presence of the *pph22*Δ*::NAT* deletion allele (Fig S3F). Specifically, NAT-sensitive progeny remained haploid, while all NAT-resistant progeny displayed peaks at 1N, 2N, and 4N, suggesting altered DNA replication or disrupted ploidy regulation in these cells. These results suggest that *PPH22* may have a role in ploidy regulation, and its loss may disrupt cell cycle regulation and lead to altered DNA inheritance patterns.

Further analysis, such as whole-genome sequencing, could help clarify which process may be maintaining *MAT* heterozygosity in the NAT-sensitive progeny. While mitochondria in the majority of progeny were inherited from the *MAT***a** parent, progeny from two out of seven basidia from independent mating spots in one of the *pph22*Δ *sup* x WT crosses inherited mitochondria from the *MAT*α parent (Fig S3E). Together, with the genotyping results from *pph22*Δ x WT crosses, these findings suggest that deletion of *PPH22* may also influence mitochondrial inheritance. Although no evidence of pseudosexual reproduction was found in the *pph22*Δ *sup* crosses, the observed α-**a** sexual reproduction was atypical, characterized by an unexpected segregation of mating-type alleles.

### Overexpression of *PPG1* suppresses loss of *PPH22*

All independently obtained *pph22*Δ *sup* mutant strains were found to be aneuploid for a segment of chromosome 6 encompassing seven genes [30]. Of these genes, *PPG1*, which encodes a putative PP2A catalytic subunit and shares significant sequence similarity with Pph22, was overexpressed 40-fold in the *pph22*Δ *sup* mutants compared to wild type [30]. To determine if *PPG1* overexpression was the causative mutation suppressing *pph22*Δ phenotypes, a *PPG1* allele with a constitutively active promoter (*P_TEF1_-PPG1-NEO)* was integrated into the safe haven 1 of the *PPH22/pph22*Δ heterozygous mutant diploid strain and homologous recombination was confirmed through PCR genotyping. Following self-filamentation on MS media, random spores were dissected and germinated on YPD (Fig 5A). Colonies containing both *NAT* and *NEO* drug resistance markers, confirmed by growth on selective media and PCR genotyping, were faster growing and larger in size than *pph22*Δ mutant colonies, suggesting that overexpression of *PPG1* can partially rescue growth defects of *pph22*Δ.

**Figure 5.**
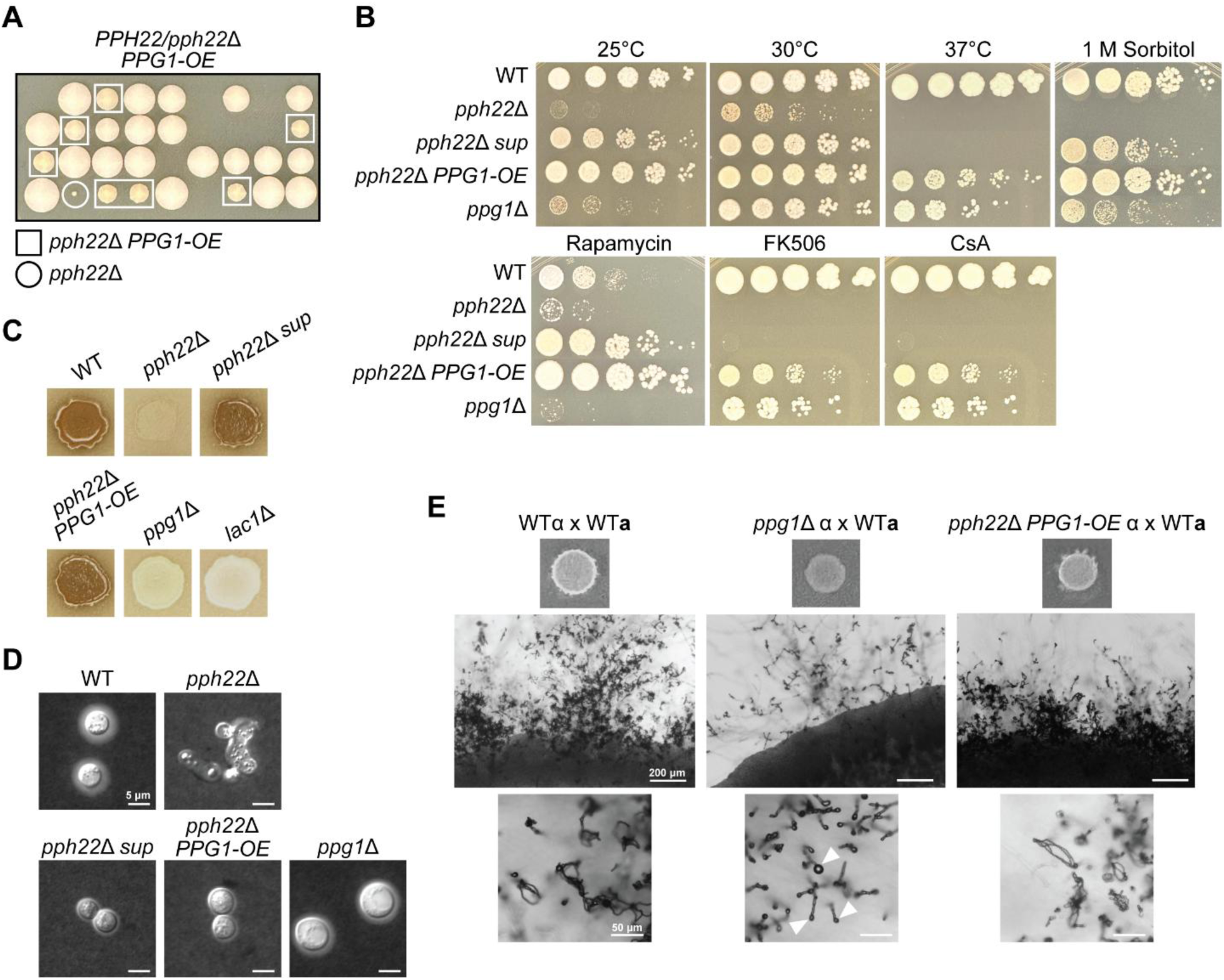
Overexpression of *PPG1* suppresses the *pph22*Δ mutation. A) Germination of spores dissected from a *PPH22/pph22*Δ heterozygous mutant diploid strain expressing *TEF1-PPG1* (*PPG1-OE)* on YPD. The genotypes of the colonies indicated were determined from growth on YPD+NAT (*pph22*Δ::*NAT* allele) and YPD+NAT+NEO (*pph22*Δ*::NAT* and *P_TEF1_-PPG1-NEO* alleles). B) The indicated strains were serially diluted and grown on YPD solid media at 25⁰C, 30⁰C, and 37⁰C; or YPD supplemented with 1 M sorbitol, 100 ng/mL rapamycin, 1 μg/mL FK506, or 100 μg/mL cyclosporine A, each at 30⁰C. Images were taken after three days incubation. C) Strains were grown in liquid culture overnight in YPD at 30⁰C and plated onto Niger seed agar. Images were captured after four days. D) Analysis of capsule formation by India ink staining. The indicated strains were grown for 3 days in RPMI media at 30⁰C. Images are representative of two biological replicates. Scale bar is 5 μm. E) Mating of *ppg1*Δ and *pph22*Δ *PPG1-OE* strains crossed to wild type on MS media. Images were taken after six weeks. White arrowheads indicate bald basidia heads that did not produce spores. The scale bars in the middle and bottom panels are 200 μm and 50 μm.

Next, to examine the role of *PPG1* overexpression in response to different nutrient and stress conditions, *pph22*Δ *PPG1-OE*, *pph22*Δ, *pph22*Δ *sup, ppg1*Δ, and wild type strains were serially diluted and spotted onto YPD at 25⁰C, 30⁰C, and 37⁰C; YPD with 1 M sorbitol; YPD with rapamycin; and YPD with the immunosuppressive drugs FK506 or cyclosporine A (CsA) (Fig 5B). *pph22*Δ exhibited severe growth defects or no growth on YPD at 25⁰C, 30⁰C, and 37⁰C, as well as on YPD supplemented with sorbitol, FK506, or cyclosporine A, which was partially reversed in the *pph22*Δ *sup* strain, consistent with previously published results. On YPD supplemented with rapamycin, on which *pph22*Δ strains do not have noticeable defects compared to wild type, both *pph22*Δ *sup* and *pph22*Δ *PPG1-OE* strains displayed significant growth advantages. Collectively, overexpression of *PPG1* in the *pph22*Δ mutant was able to fully restore or nearly restore growth to wild-type levels under each condition tested, suggesting that *P_TEF1_-PPG1* is more effective at compensating for *pph22*Δ than overexpression of *PPG1* due to aneuploidy in *pph22*Δ *sup* strains.

It was surprising that *ppg1*Δ did not phenocopy *pph22*Δ, and only exhibited growth defects on YPD at 25⁰C and on YPD supplemented with sorbitol or rapamycin, compared to growth on YPD at 30⁰C. In fact, *ppg1*Δ displayed similar growth to *pph22*Δ *PPG1-OE* under several conditions, including growth at 30⁰C and 37⁰C as well as in the presence of FK506 and CsA. These results suggest that while Pph22 and Ppg1 are hypothesized to have similar functions and may play partially compensatory roles, Pph22 is likely the primary catalytic subunit of PP2A, as its loss leads to more severe effects than the loss of *PPG1.* To assess the abilities of *ppg1*Δ and *pph22*Δ *PPG1-OE* strains to produce melanin pigment and polysaccharide capsule, important virulence factors to protect cells from the host immune response during *Cryptococcus* infection. To induce melanin and capsule production, cells were grown on Niger seed agar and in RPMI media, respectively, with both incubated at 30⁰C.

*pph22*Δ *sup* and *pph22*Δ *PPG1-OE* produced similar amounts of melanin and were comparable to the wild type, while *ppg1*Δ could not produce melanin, similar to the *lac1*Δ negative control (Fig 5C). These results suggest that *PPG1* is required for melanin production, while *PPG1* overexpression in the *pph22*Δ background is sufficient to restore this virulence trait. India ink counterstaining of RPMI-grown cells showed that *PPG1* overexpression restored normal cell morphology in the *pph22*Δ mutant, though capsule production was still impaired (Fig 5D). In contrast, *ppg1*Δ cells could produce capsule, albeit to a lesser extent than wild-type cells. Together, these findings are consistent with the idea that *PPG1* can functionally compensate for the loss of *PPH22* in certain contexts.

To further explore the functional differences between Pph22 and Ppg1, 3D structural models of the STRIPAK complex with either Pph22 or Ppg1 as the catalytic subunit were generated with AlphaFold3 [50]. The resulting Pph22-containing structure was comparable to the previously published model generated with AlphaFold2, though slight differences in folding were observed [30, 51]. The two models appeared largely similar, indicating that Ppg1 can potentially assemble into a complex with the other STRIPAK subunits (Fig S4A, S4B). Cryo-EM studies of the human STRIPAK complex have demonstrated that Far8 forms a homotetramer, serving as a scaffold at the base of the complex [12]. To assess whether *Cryptococcus* Far8 could similarly assemble into a tetramer, a structural model was generated using four copies of the Far8 protein (Fig S4C). This model was then combined with either the Pph22-Tpd3 or Ppg1-Tpd3 heterodimers to examine PP2A assembly. Both models appeared highly similar overall, though the four Far8 proteins adopted slightly different conformations in each structure (Fig S4D, S4E). These findings suggest that both Pph22 and Ppg1 can potentially integrate into STRIPAK in a comparable manner, perhaps filling analogous roles within the complex.

Finally, the involvement of Ppg1 during mating was examined by crossing *ppg1*Δ and *pph22*Δ *PPG1-OE* strains with wild type. Similar to what has been observed in *pph22*Δ crosses, *ppg1*Δ x WT exhibited delayed mating and hyphae were produced only in isolated spots on the mating patch after six weeks of incubation (Fig 5E compared to Fig 1A). With further incubation, more extended hyphal branches were produced but mostly bald basidia heads with almost no spore formation. In contrast, *pph22*Δ *PPG1-OE* x WT crosses displayed signs of mating at a similar rate to wild type and produced abundant basidia with spore chains (Fig 5E compared to Fig 4A). These findings indicate that Ppg1 plays a crucial role during mating and sexual development, similar to Pph22. Taken together, this further supports the conclusion that *PPG1* overexpression is the key mutation suppressing *pph22*Δ, as targeted overexpression of *PPG1* in the *pph22*Δ background similarly restored both mating and growth deficiencies.

### Pseudosexual reproduction also occurs at a high rate in *far8*Δ mutant crosses

It has been reported that pseudosexual reproduction occurs at a rate of ∼1% between wild-type mating partners with compatible parental genomes, as well as in partners with incompatible parental genomes due to genetic divergence or genome structure variation [38, 52]. The analyses presented here demonstrated that deletion of *PPH22* results in exclusively pseudosexual reproduction following α-**a** sexual mating. To further investigate whether other mutations in the STRIPAK complex influence pseudosexual reproduction, *far8*Δ mutant crosses were analyzed. Far8 has also been shown to be critical during *Cryptococcus* sexual development, with *far8*Δ strains exhibiting severe defects in hyphal production and sporulation. To inspect the sexual structures of *far8*Δ x WT crosses in higher resolution, hyphae, basidia, and spore chains were imaged with SEM (Fig 6A). Spores were observed budding from the basidia in irregular numbers, often exceeding the typical set of four. In some cases, seven or more spores emerged independently from separate sites on a single basidium. *far8*Δ strains have been predominantly identified as diploid, suggesting that mating could result in triploid meiosis. This raises the possibility that the observed irregular spore structures may be influenced by the unique genomic composition in the basidium.

**Figure 6.**
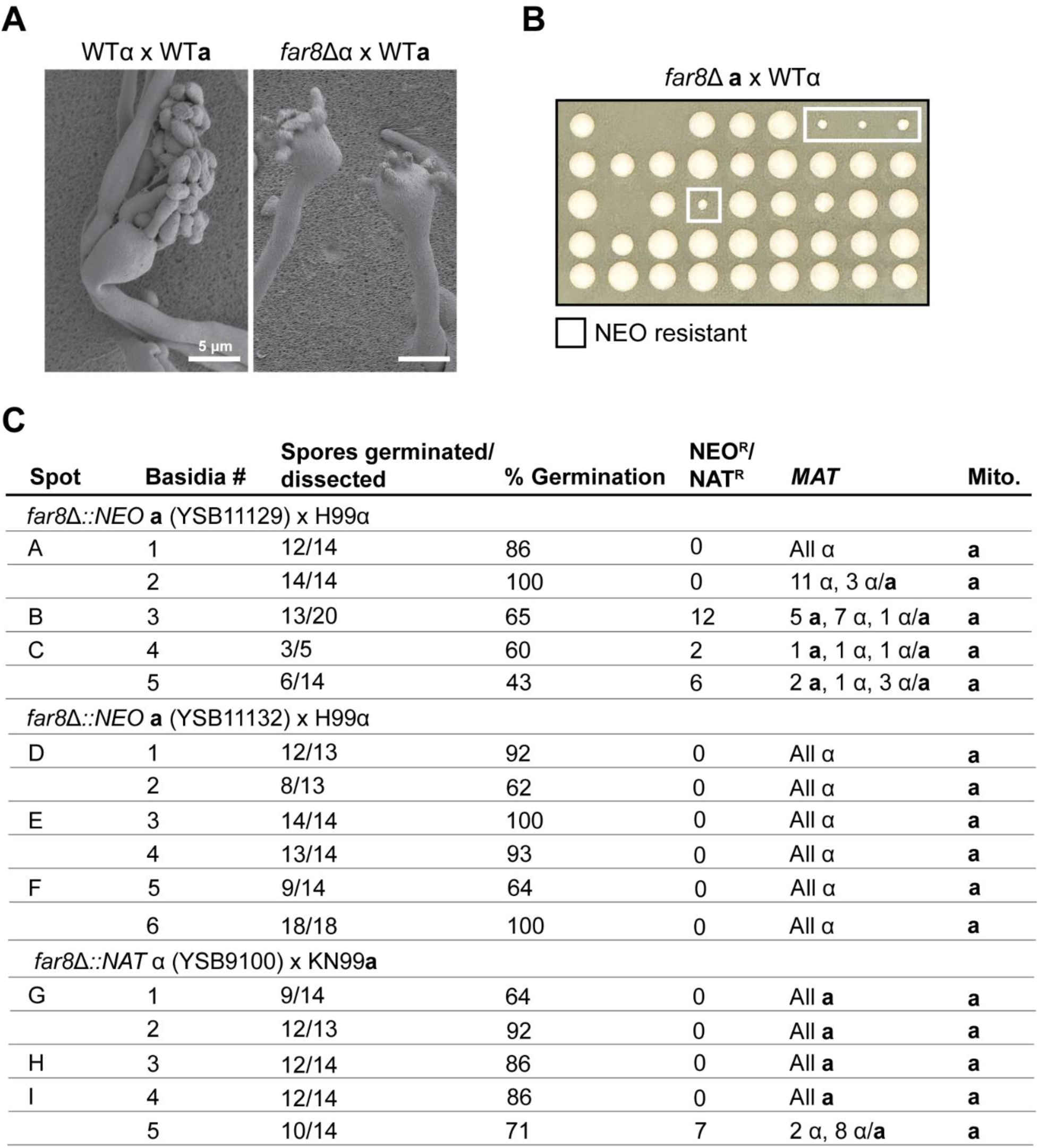
*far8*Δ mutants also display a high rate of pseudosexual reproduction. A) SEM of mated wild-type and *far8*Δ shows two basidia heads producing spores with abnormal morphology. In the basidium on the right, spores are seen budding from seven independent spots on the basidial head. The scale bar represents 5 μm. B) Dissection of spores from *far8*Δ x WT yields colonies mostly homogenous in size on YPD. *far8*Δ mutants were identified based on growth on YPD+NEO and form smaller, slow-growing colonies compared to wild type. Each row of spores is from an independent basidium. C) Genotyping analysis of progeny from individual basidia in *far8*Δ x WT crosses. Each spot analyzed is from an independent mating experiment.

Next, spores were dissected from individual basidia from both *far8*Δ **a** x WTα and *far8*Δ α x WT**a** crosses. Spores that germinated formed mostly homogenously sized colonies on YPD and did not contain the drug resistance marker, but occasionally small, slow-growing *far8*Δ mutant colonies were obtained (Fig 6B). In total, spores from 16 basidia obtained from nine independent mating spots were dissected and genotyped (Fig 6C). Given the genomic instability of triploid meiosis, reduced spore viability was expected. However, surprisingly, 10 out of 16 basidia exhibited germination rates above 70%, with an overall average of 79%, which is comparable to the germination rate observed in progeny from wild-type crosses. PCR genotyping of the *MAT* locus and mitochondrial DNA in progeny from 11 out of 16 dissected basidia were consistent with pseudosexual reproduction, characterized by uniparental nuclear inheritance from only the wild-type parent, a pattern also observed in *pph22*Δ mutant crosses, and mitochondrial inheritance from the *MAT***a** parent. Heterozygosity at the *MAT* locus was also detected in some progeny from the remaining five basidia of the *far8*Δ x WT crosses, resembling the findings in the *pph22*Δ *sup* x WT crosses. Notably, 11 out of 16 progeny possessing both *MAT***a** and *MAT*α also had the *far8*Δ mutant allele, which may be attributed to the altered genomic structure of the *far8*Δ parental strains. Taken together, these results suggest that loss of STRIPAK components *PPH22* or *FAR8* drives pseudosexual reproduction, with the mutant nuclei at a selective disadvantage to the wild-type nucleus, as shown by uniparental nuclear inheritance predominantly from the wild-type parent across multiple independent crosses (Fig S5).

## Discussion

The STRIPAK complex has been broadly recognized across fungi as a critical regulator of both asexual and sexual development. In *C. neoformans*, we previously demonstrated that mutants defective in STRIPAK components exhibit defects in several developmental processes, along with increased genome instability, leading to segmental and whole-chromosome duplications or losses [30]. Additionally, STRIPAK mutations were shown to impact the production of virulence traits and pathogenesis during infection. In this study, we further explored the specific roles of STRIPAK during mating and sexual development, identifying two genes, *PPH22* and *FAR8*, that were directly involved in pseudosexual reproduction. This unusual reproductive strategy, which was previously characterized in our lab, occurs when there is incompatibility between mating partners due to genetic divergence or genome structure variation, allowing *C. neoformans* to overcome reproductive barriers [38]. While previously predicted to occur at low frequencies, pseudosexual reproduction enables the production of viable offspring in conditions where successful α-**a** sexual reproduction is not possible or is inefficient, contributing to survival and adaptation.

It has been hypothesized that pseudosexual reproduction may preferentially favor the predominance of one mating type over the other and that specific factors influence which nucleus is selectively lost following mating. Our findings provide evidence for genetic factors influencing pseudosexual reproduction, as progeny from crosses involving *pph22*Δ or *far8*Δ mutants exclusively inherited their nuclear genome from the wild-type parent. This supports the hypothesis that, even in cases where mating partners are genetically compatible, intrinsic gene variation can endow one genome with a selective advantage over the other. Such selective inheritance could potentially contribute to fungal adaptation, as it might allow for the retention of beneficial genetic traits while discarding less advantageous ones. Given that loss of *PPH22* or *FAR8* leads to widespread genome instability, including segmental and whole chromosomal aneuploidy, it is likely that these defects impose a fitness cost during sexual development, favoring the retention of the more stable wild-type genome. The defective spore formation observed in *pph22*Δ and *far8*Δ mutants, along with the presence of aberrant DNA content in the progeny, suggests that these mutations may impair proper chromosome segregation or meiotic progression. As a result, selective nuclear retention in pseudosexual reproduction may serve as a mechanism to safeguard genomic integrity, ensuring that only nuclei with fewer deleterious mutations or with more favorable genomic configurations contribute to the next generation.

During *Cryptococcus* sexual development, the process of nuclear migration, segregation of nuclei within dikaryotic hyphae, and meiosis are tightly coordinated processes that ensure proper exchange and inheritance of genetic material [34, 53]. STRIPAK has been implicated in actin cytoskeletal organization in other organisms, raising the possibility that its components may play a role in regulating nuclear dynamics during mating [18, 54–57]. In *Cryptococcus*, sexual reproduction involves an extended dikaryotic phase prior to karyogamy, requiring precise nuclear positioning within hyphae. Our fluorescence microscopy data indicate that *pph22*Δ mutants exhibit nuclear segregation defects, as their nuclei frequently fail to reach the basidium. This suggests that *PPH22* may be required for proper nuclear migration during hyphal branching and elongation. However, the observation that wild-type nuclei were sometimes lost without generating viable *pph22*Δ spores suggests that *PPH22* also plays a direct role in meiosis. These findings highlight the role of STRIPAK components in maintaining nuclear integrity throughout sexual development.

Interestingly, although recessive mutations are typically complemented in a dikaryon by the presence of a wild-type allele in the opposing nucleus, these findings provide additional evidence that *pph22* mutations are haploinsufficient in this context. Developmental defects were observed in dikarya formed between *pph22*Δ and wild-type strains, and importantly, it was previously demonstrated that a *PPH22/pph22*Δ heterozygous diploid is haploinsufficient, supporting the conclusion that a single copy of *PPH22* is insufficient to support normal development in a diploid or dikaryon. This dosage sensitivity may in part explain the dominance of the wild-type genome in pseudosexual reproduction. Additionally, frequent aneuploidy observed in the *pph22*Δ mutant strains may further compromise nuclear division and migration or meiotic progression, contributing to the nuclear selection bias. Together, these data suggest that both haploinsufficiency and genomic instability may underlie the strong inheritance bias and disrupted sexual development seen in *pph22*Δ crosses, potentially serving as a mechanism to eliminate defective nuclei and maintain genome integrity.

The relationship between *PPH22* and *PPG1* suggests functional redundancy, as overexpression of *PPG1* can compensate for *pph22*Δ during both vegetative growth and sexual development. This compensation indicates that *PPG1* can partially fulfill the role of *PPH22* in regulating key cellular processes, likely due to their structural and functional similarities.

However, despite this redundancy, *PPH22* appears to have a more critical role, as *ppg1*Δ mutants exhibited more subtle phenotypes compared to the severe defects observed in *pph22*Δ. Notably, while deletion of *PPH22* is synthetically lethal with calcineurin inhibition by FK506 or cyclosporine A, deletion of *PPG1* did not cause a significant growth defect under these conditions. This suggests that *PPH22* plays a predominant role in maintaining essential phosphatase activity that becomes indispensable when calcineurin is inhibited, whereas *PPG1* may function as a secondary phosphatase with limited capacity to support survival under these stresses. These findings reinforce the idea that *PPH22* is the primary catalytic subunit of PP2A, with *PPG1* providing only partial redundancy. This difference in phenotypic severity highlights the distinct regulatory contributions of these phosphatases, with *PPH22* playing a dominant role in STRIPAK-dependent processes.

Overall, our findings highlight the critical role of the STRIPAK complex in coordinating key aspects of *Cryptococcus* sexual reproduction and development, including nuclear dynamics during mating, genome stability, and meiotic progression. While STRIPAK has been implicated in developmental processes across diverse fungal species, its role in pseudosexual reproduction is intriguing and unique to *Cryptococcus,* where it seems to bear a resemblance to hybridogenesis observed in some animal species. The ability to selectively retain one parental nucleus during mating may serve as a mechanism to maintain genome integrity in the face of genetic instability, ensuring the production of viable progeny when conventional mating pathways are disrupted. Our study provides new insights into the genetic factors that govern sexual reproduction in *Cryptococcus*, shedding light on how STRIPAK complex mutations influence nuclear selection and inheritance. Future studies will explore how STRIPAK coordinates with other signaling pathways, such as calcineurin and MAPK regulation, to direct cellular growth processes. Additionally, investigating whether similar mechanisms operate in other fungal pathogens could reveal broader implications for genome evolution and adaptation.

## Materials and methods

### Strains, media, and growth conditions

Strains characterized in this study are listed in Table S1. Strains were prepared for long-term storage as 20% glycerol stocks at −80°C. Fresh cultures were revived and maintained on YPD (1% yeast extract, 2% Bacto Peptone, 2% dextrose) agar medium, incubated at 30°C. *Cryptococcus* transformants harboring dominant drug-resistance markers were selected on YPD medium supplemented with 100 μg/mL nourseothricin (NAT), 200 μg/mL neomycin (G418), or 100 μg/mL hygromycin B (HYG). Strains were grown in YPD liquid cultures at 30°C as indicated. For plate growth assays, strains were cultivated on YPD, Murashige and Skoog (MS) (Sigma-Aldrich M5519), or Niger seed (7% Niger seed, 0.1% dextrose). Sorbitol was added to YPD medium at a 1 M concentration. To analyze cell growth in response to immunosuppressive agents, rapamycin (100 ng/mL), FK506 (1 μg/mL), and cyclosporine A (100 μg/mL) were added to YPD medium. For serial dilution assays, fresh cells were diluted to a starting OD_600_ of 0.1, serially diluted 5-fold, and spotted onto plates for the indicated media and temperature conditions. Plates were incubated for two to seven days and photographed daily. For capsule analysis, strains were incubated for 3 days in RPMI media at 30°C, followed by negative staining with India ink. For mating assays, strains were incubated on MS plates, face up, in the dark at room temperature for up to eight weeks. Spores dissected from MS plates were germinated on YPD at 30°C for five days.

### Strain construction

Mutant strains analyzed in this study were generated in the *C. neoformans* H99α, YL99**a,** or KN99**a**/KN99α backgrounds. All genetic manipulations were done using CRISPR-Cas9-mediated mutagenesis and the TRACE system, according to previously published protocols [58, 59]. In brief, Cas9 was amplified from the pBHM2403 plasmid and gRNAs were generated with sequence specific-primers and amplification of the gRNA scaffold from the pBHM2329 plasmid [59]. Donor DNA templates for transformations were generated by PCR or plasmid digestion and purified with the QIAquick gel extraction kit (Qiagen #28704). Cells and DNA were prepared for transformation as described in the TRACE system, and electroporated utilizing a BioRad gene pulser (0.45 kV, 125 μF, 600 Ω). Cells were recovered in YPD media for 2-3 hours at 30⁰C following electroporation.

To generate the *PPG1* overexpression allele, the *TEF1* promoter (1.2 kb) and *PPG1* coding regions were amplified from H99α genomic DNA and assembled into plasmid pSDMA57, digested with XhoI and SpeI, via Gibson assembly (New England Biolabs E5510S) for targeting to the safe haven (SH) locus [49]. GFP tagging of Nop1 was also performed through integration at the safe haven. To facilitate homologous recombination at the target locus, the safe haven plasmid pYSCE5 (pSH-P_H3_-GFP-HYG) was designed to include a safe haven genomic sequence containing AscI and PacI restriction sites for linearization, along with the histone H3 promoter, GFP, and the hygromycin B resistance gene (HYG). The *NOP1* gene, amplified from H99α genomic DNA without its stop codon, was incorporated into plasmid pYSCE5 between the *H3* promoter and GFP, generating pYSCE8 (pSH-H3-NOP1-GFP-HYG). Plasmids for targeted integration at the safe haven were linearized via overnight digestion with AscI and PacI and gel-purified prior to transformation. The *ppg1::NEO* deletion allele was made via PCR amplification of approximately 1 kb homologous 5’ and 3’ flanking regions to the *PPG1* open reading frame, which were then assembled with the neomycin resistance gene expression cassette (NEO) through overlap PCR [60, 61]. Transformants were plated onto selective media (YPD+NEO or YPD+HYG) and allowed to grow up to 7 days at 30⁰C. Stable genomic integrants were verified by diagnostic PCR amplifying the 5′ and 3′ flanking junctions of the insertion site.

### Primers, PCR genotyping, and RFLP analysis

Primers employed in this study are listed in Table S2. Genomic DNA from strains generated in this study was extracted from cells with the MasterPure Yeast DNA Purification Kit (LGC Biosearch Technologies). Dissected progeny from mated crosses were genotyped by colony PCR. Single colonies were picked into 0.25% SDS and 1 mM EDTA, briefly frozen at - 20 °C, and then heated at 100⁰C for 10 minutes in a thermocycler. The resulting supernatants were used as DNA templates for PCR. To determine the mating type of *C. neoformans* strains, the *STE20* gene in the *MAT* locus was amplified using primers specific to the *STE20*α and *STE20***a** alleles. The mitochondrial *COX1* gene, which differs in size in KN99 background strains compared to H99 due to the presence of an intron, was probed to determine mitochondrial inheritance. For KN99α *pph22*Δ *sup* crosses, the KN99**a** SSH118 strain, which has a recombinant mitochondrial genome, was used as the wild-type parent to differentiate between mitochondrial genotypes in the KN99 background progeny. After PCR amplification of *COX1*, the resulting PCR product was digested with restriction enzyme BsrI for restriction fragment length polymorphism (RFLP) analysis. Digestion of the *COX1* PCR product amplified from KN99**a**, which contains three BsrI recognition sites, and SSH118, which has two BsrI sites, were used as controls to determine the mitochondrial inheritance of the progeny from *pph22*Δ *sup* x SSH118 crosses.

### Microscopy

For sample preparation for scanning electron microscopy from mated strains of *Cryptococcus*, an agar slice of the plated cells was fixed in a solution of 4% formaldehyde and 4% glutaraldehyde for 2 hours at 4°C. The fixed cells were then gradually dehydrated in a graded ethanol series (30%, 50%, 70%, and 95%), with a 15-minute incubation at 4°C for each concentration. This was followed by three washes with 100% ethanol, each for 15 minutes at 4°C. The samples were further dehydrated using a Ladd CPD3 Critical Point Dryer and coated with a layer of gold using a Denton Desk V Sputter Coater (Denton Vacuum, USA). Hyphae, basidia, and basidiospores were observed with a scanning electron microscope with an EDS detector (Apreo S, ThermoFisher, USA). Brightfield, differential interference contrast (DIC), and fluorescence microscopy images were visualized with an AxioScop 2 fluorescence microscope and captured with an AxioCam MRm digital camera (Zeiss, Germany). Consistent exposure times were used for all images captured from the same experiment. Image processing and merging channels of fluorescent images was performed with ImageJ/Fiji software.

### Cell-cell fusion and mating assays

To induce mating, the indicated strains were grown overnight in YPD medium, diluted to an OD_600_ of 1.0, and 4 μL of each cell suspension was spotted onto MS agar plates. The plates were incubated at room temperature in the dark and monitored for signs of filamentation and sporulation for up to eight weeks. For wild-type mating crosses, *MAT***a** cells (KN99**a**) were mixed with *MAT*α cells (H99α) in equal amounts and spotted onto MS plates. For crosses involving *pph22*Δ and *far8*Δ strains, the following modifications were made to compensate for the impaired growth of these mutants on MS medium. Fresh *pph22*Δ and *far8*Δ mutant cells were first spotted onto MS agar and pre-incubated in the dark at room temperature for two days, without the wild-type mating partner. After pre-growth, wild-type cells of the opposite mating type were added directly on top of the mutant cells at a ten-fold lower cell amount. Twelve mating spots were prepared per plate, with each spotted treated as an independent replicate. A wild-type H99α x KN99**a** cross was included on each plate as a control. Basidiospores were dissected from individual basidia, or randomly dissected, and incubated on YPD medium at 30⁰C for 3 to 4 days to determine spore viability and germination rates.

To assess cell-cell fusion frequency in wild type x *pph22*Δ crosses, KN99**a** or KN99α strains carrying a NEO resistance marker (JOHE10493, JOHE18842) were mixed in 1:10 proportions with *pph22*Δ*:NAT* **a** or α strains, spotted onto MS agar, and incubated in the dark at room temperature for 3 days. A KN99**a***::NAT* (JOHE18853) x KN99α*::NEO* (JOHE18842) wild-type cross was used as a control. The entire patch of cells was harvested and serially diluted in PBS before plating onto YPD+NAT, YPD+NEO, and YPD+NAT+NEO selective media. Colony-forming units (CFUs) were counted after incubation at 30°C for two days (for wild-type selection) or four days (for pph22Δ and dual-resistant fusion products). Fusion frequency was calculated as the number of CFUs on YPD+NAT+NEO divided by the total number of colonies on YPD+NAT and YPD+NEO, and expressed as a percentage relative to the wild-type control. Each assay was performed in triplicate.

### Whole-genome sequencing and SNP analysis

Genomic DNA for whole-genome sequencing was extracted from saturated 4 mL YPD cultures with the MasterPure Yeast DNA Purification Kit. The precipitated DNA was dissolved in 35 μL of 1x TE buffer (100 mM Tris-HCl, 10 mM EDTA, pH=8.0), and the concentration was estimated using Qubit. Illumina sequencing was performed at the Duke Sequencing and Genomic Technologies core facility (https://genome.duke.edu) with Novaseq X Plus, providing 250 bp paired-end reads. The Illumina sequences were trimmed using trim-galore (v 0.6.7) and mapped to the H99α genome assembly (RefSeq: GCF_000149245.1) using Geneious software. The resulting BAM files were converted to TDF format, and read coverage was visualized in IGV to verify uniform read coverage across the genome. For SNP calling, the Illumina sequences were mapped to the H99α genome assembly using the Geneious default mapper with five iterations. Variant calling was performed with BAM mapped read files, with parameters set to a 0.9 variant frequency and a minimum of 100x coverage per variant. For the KN99**a**/KN99α diploid strain (CnLC6683), variant calling was performed with a 0.45 frequency and 100x coverage per variant. Illumina sequences from H99α and KN99**a**/KN99α served as controls for SNP calling analysis. Variants were exported from Geneious as VCF files and imported into IGV for visualization.

### Flow cytometry

For ploidy determination of *Cryptococcus* samples, fluorescence activated cell sorting (FACS) was performed according to a previously published protocol, with slight modifications [62]. Wild-type and mutant strains were grown on YPD plate medium at 30⁰C overnight, harvested, and washed with PBS. The cells were then fixed in 70% ethanol for 16 hours at 4⁰C. Fixed cells were pelleted and washed with 1 mL NS buffer (10 mM Tris-HCl, 0.25 M sucrose, 1 mM EDTA, 1 mM MgCl_2_, 0.1 mM ZnCl_2_, 0.4 mM phenylmethylsulfonyl fluoride, and 7 mM β-mercaptoethanol). After centrifugation, the cells were treated with RNase (0.5 mg/mL) and stained with propidium iodide (10 μg/mL) in a 200 μL suspension of NS buffer for 2 hours in the dark. Then, 50 μL of the stained cells were diluted into 200 μL of 50 mM Tris-HCl, pH=8.0, and submitted to the Duke Cancer Institute Flow Cytometry Shared Resource for analysis. Fluorescence was measured using a BD FACSCanto flow cytometer and analyzed with BD FACSDiva software. Approximately 15,000 to 20,000 events were analyzed for each sample.

### Quantification and statistical analysis

All statistical analyses and data visualizations were performed with GraphPad Prism v10 software. Details regarding replication, quantification, and statistical tests used for each experiment are provided in the methods and corresponding figure legends.

## Acknowledgements

PPP is supported by NIH/NIAID T32 grant AI052080-21 as a Tri-I MMPTP fellow. This work is also supported by NIH/NIAID R01 grants AI039115-27, AI050113-20, and AI172451-03 awarded to JH. JH is a co-director of the CIFAR Fungal Kingdom: Threats & Opportunities program. We thank Dr. Vikas Yadav for guidance and expertise and our laboratory manager Anna Floyd Averette for technical support.

## References

1. Goudreault M, D’Ambrosio LM, Kean MJ, Mullin MJ, Larsen BG, Sanchez A, et al. A PP2A phosphatase high density interaction network identifies a novel striatin-interacting phosphatase and kinase complex linked to the cerebral cavernous malformation 3 (CCM3) protein. Mol Cell Proteomics. 2009;8(1):157–71. Epub 2008/09/11. doi: 10.1074/mcp.M800266-MCP200. PubMed PMID: 18782753; PubMed Central PMCID: 2621004.

2. Neisch AL, Neufeld TP, Hays TS. A STRIPAK complex mediates axonal transport of autophagosomes and dense core vesicles through PP2A regulation. J Cell Biol. 2017;216(2):441–61. Epub 2017/01/20. doi: 10.1083/jcb.201606082. PubMed PMID: 28100687; PubMed Central PMCID: PMC5294782.

3. Li D, Musante V, Zhou W, Picciotto MR, Nairn AC. Striatin-1 is a B subunit of protein phosphatase PP2A that regulates dendritic arborization and spine development in striatal neurons. J Biol Chem. 2018;293(28):11179–94. Epub 2018/05/29. doi: 10.1074/jbc.RA117.001519. PubMed PMID: 29802198; PubMed Central PMCID: PMC6052221.

4. Gordon J, Hwang J, Carrier KJ, Jones CA, Kern QL, Moreno CS, et al. Protein phosphatase 2a (PP2A) binds within the oligomerization domain of striatin and regulates the phosphorylation and activation of the mammalian Ste20-Like kinase Mst3. BMC Biochemistry. 2011;12:54. doi: 10.1186/1471-2091-12-54. PubMed PMID: 21985334; PubMed Central PMCID: 3217859.

5. Janssens V, Goris J. Protein phosphatase 2A: a highly regulated family of serine/threonine phosphatases implicated in cell growth and signalling. Biochem J. 2001;353(Pt 3):417–39. doi: 10.1042/0264-6021:3530417. PubMed PMID: 11171037; PubMed Central PMCID: PMC1221586.

6. Cho US, Xu W. Crystal structure of a protein phosphatase 2A heterotrimeric holoenzyme. Nature. 2007;445(7123):53-7. Epub 20061101. doi: 10.1038/nature05351. PubMed PMID: 17086192.

7. Sandal P, Jong CJ, Merrill RA, Song J, Strack S. Protein phosphatase 2A - structure, function and role in neurodevelopmental disorders. J Cell Sci. 2021;134(13). Epub 20210706. doi: 10.1242/jcs.248187. PubMed PMID: 34228795; PubMed Central PMCID: PMC8277144.

8. Sents W, Ivanova E, Lambrecht C, Haesen D, Janssens V. The biogenesis of active protein phosphatase 2A holoenzymes: a tightly regulated process creating phosphatase specificity. FEBS J. 2013;280(2):644–61. Epub 20120425. doi: 10.1111/j.1742-4658.2012.08579.x. PubMed PMID: 22443683.

9. Ribeiro PS, Josue F, Wepf A, Wehr MC, Rinner O, Kelly G, et al. Combined functional genomic and proteomic approaches identify a PP2A complex as a negative regulator of Hippo signaling. Mol Cell. 2010;39(4):521–34. Epub 2010/08/28. doi: 10.1016/j.molcel.2010.08.002. PubMed PMID: 20797625.

10. Kuck U, Radchenko D, Teichert I. STRIPAK, a highly conserved signaling complex, controls multiple eukaryotic cellular and developmental processes and is linked with human diseases. Biol Chem. 2019. Epub 2019/05/03. doi: 10.1515/hsz-2019-0173. PubMed PMID: 31042639.

11. Kuck U, Poggeler S. STRIPAK, a fundamental signaling hub of eukaryotic development. Microbiol Mol Biol Rev. 2024;88(4):e0020523. Epub 20241111. doi: 10.1128/mmbr.00205-23. PubMed PMID: 39526753; PubMed Central PMCID: PMC11653735.

12. Jeong BC, Bae SJ, Ni L, Zhang X, Bai XC, Luo X. Cryo-EM structure of the Hippo signaling integrator human STRIPAK. Nature Structural & Molecular Biology. 2021;28(3):290-9 Epub 2021/02/27. doi: 10.1038/s41594-021-00564-y. PubMed PMID: 33633399; PubMed Central PMCID: PMC8315899.

13. Shi Z, Jiao S, Zhou Z. STRIPAK complexes in cell signaling and cancer. Oncogene. 2016;35(35):4549–57. Epub 2016/02/16. doi: 10.1038/onc.2016.9. PubMed PMID: 26876214.

14. Bernhards Y, Poggeler S. The phocein homologue SmMOB3 is essential for vegetative cell fusion and sexual development in the filamentous ascomycete *Sordaria macrospora*. Curr Genet. 2011;57(2):133–49. Epub 20110113. doi: 10.1007/s00294-010-0333-z. PubMed PMID: 21229248; PubMed Central PMCID: PMC3059760.

15. Bloemendal S, Bernhards Y, Bartho K, Dettmann A, Voigt O, Teichert I, et al. A homologue of the human STRIPAK complex controls sexual development in fungi. Mol Microbiol. 2012;84(2):310–23. doi: 10.1111/j.1365-2958.2012.08024.x. PubMed PMID: 22375702.

16. Kuck U, Stein V. STRIPAK, a Key Regulator of Fungal Development, Operates as a Multifunctional Signaling Hub. J Fungi. 2021;7(6):443. Epub 2021/07/03. doi: 10.3390/jof7060443. PubMed PMID: 34206073; PubMed Central PMCID: PMC8226480.

17. Chen A, Liu N, Xu C, Wu S, Liu C, Qi H, et al. The STRIPAK complex orchestrates cell wall integrity signalling to govern the fungal development and virulence of *Fusarium graminearum*. Mol Plant Pathol. 2023;24(9):1139–53. Epub 20230606. doi: 10.1111/mpp.13359. PubMed PMID: 37278525; PubMed Central PMCID: PMC10423325.

18. Pracheil T, Thornton J, Liu Z. TORC2 signaling is antagonized by protein phosphatase 2A and the Far complex in *Saccharomyces cerevisiae*. Genetics. 2012;190(4):1325–39. doi: 10.1534/genetics.111.138305. PubMed PMID: 22298706; PubMed Central PMCID: 3316646.

19. Pracheil T, Liu Z. Tiered assembly of the yeast Far3-7-8-9-10-11 complex at the endoplasmic reticulum. J Biol Chem. 2013;288(23):16986–97. doi: 10.1074/jbc.M113.451674. PubMed PMID: 23625923; PubMed Central PMCID: 3675630.

20. Frost A, Elgort MG, Brandman O, Ives C, Collins SR, Miller-Vedam L, et al. Functional repurposing revealed by comparing *S. pombe* and *S. cerevisiae* genetic interactions. Cell. 2012;149(6):1339–52. doi: 10.1016/j.cell.2012.04.028. PubMed PMID: 22682253.

21. Poggeler S, Kuck U. A WD40 repeat protein regulates fungal cell differentiation and can be replaced functionally by the mammalian homologue striatin. Eukaryot Cell. 2004;3(1):232–40. doi: 10.1128/EC.3.1.232-240.2004. PubMed PMID: 14871953; PubMed Central PMCID: PMC329509.

22. Dettmann A, Heilig Y, Ludwig S, Schmitt K, Illgen J, Fleissner A, et al. HAM-2 and HAM-3 are central for the assembly of the *Neurospora* STRIPAK complex at the nuclear envelope and regulate nuclear accumulation of the MAP kinase MAK-1 in a MAK-2-dependent manner. Mol Microbiol. 2013;90(4):796–812. Epub 2013/09/14. doi: 10.1111/mmi.12399. PubMed PMID: 24028079.

23. Wang CL, Shim WB, Shaw BD. *Aspergillus nidulans* striatin (StrA) mediates sexual development and localizes to the endoplasmic reticulum. Fungal Genet Biol. 2010;47(10):789–99. Epub 20100626. doi: 10.1016/j.fgb.2010.06.007. PubMed PMID: 20601045.

24. Elramli N, Karahoda B, Sarikaya-Bayram O, Frawley D, Ulas M, Oakley CE, et al. Assembly of a heptameric STRIPAK complex is required for coordination of light-dependent multicellular fungal development with secondary metabolism in *Aspergillus nidulans*. PLoS Genet. 2019;15(3):e1008053. Epub 2019/03/19. doi: 10.1371/journal.pgen.1008053. PubMed PMID: 30883543; PubMed Central PMCID: PMC6438568.

25. Islam KT, Bond JP, Fakhoury AM. FvSTR1, a striatin orthologue in *Fusarium virguliforme*, is required for asexual development and virulence. Appl Microbiol Biotechnol. 2017;101(16):6431–45. Epub 2017/06/24. doi: 10.1007/s00253-017-8387-1. PubMed PMID: 28643182.

26. Yamamura Y, Shim WB. The coiled-coil protein-binding motif in *Fusarium verticillioides* Fsr1 is essential for maize stalk rot virulence. Microbiology. 2008;154(Pt 6):1637–45. doi: 10.1099/mic.0.2008/016782-0. PubMed PMID: 18524918.

27. Wang CL, Shim WB, Shaw BD. The *Colletotrichum graminicola* striatin orthologue Str1 is necessary for anastomosis and is a virulence factor. Mol Plant Pathol. 2016;17(6):931–42. Epub 20160218. doi: 10.1111/mpp.12339. PubMed PMID: 26576029; PubMed Central PMCID: PMC6638439.

28. Schmidpeter J, Dahl M, Hofmann J, Koch C. ChMob2 binds to ChCbk1 and promotes virulence and conidiation of the fungal pathogen *Colletotrichum higginsianum*. BMC Microbiol. 2017;17(1):22. Epub 20170119. doi: 10.1186/s12866-017-0932-7. PubMed PMID: 28103800; PubMed Central PMCID: PMC5248491.

29. Kim MS, Zhang H, Yan H, Yoon BJ, Shim WB. Characterizing co-expression networks underpinning maize stalk rot virulence in *Fusarium verticillioides* through computational subnetwork module analyses. Sci Rep. 2018;8(1):8310. Epub 20180529. doi: 10.1038/s41598-018-26505-2. PubMed PMID: 29844502; PubMed Central PMCID: PMC5974142.

30. Peterson PP, Choi JT, Fu C, Cowen LE, Sun S, Bahn YS, et al. The *Cryptococcus neoformans* STRIPAK complex controls genome stability, sexual development, and virulence. PLoS Pathog. 2024;20(11):e1012735. Epub 20241119. doi: 10.1371/journal.ppat.1012735. PubMed PMID: 39561188; PubMed Central PMCID: PMC11614259.

31. Giles SS, Dagenais TR, Botts MR, Keller NP, Hull CM. Elucidating the pathogenesis of spores from the human fungal pathogen *Cryptococcus neoformans*. Infect Immun. 2009;77(8):3491–500. Epub 20090518. doi: 10.1128/IAI.00334-09. PubMed PMID: 19451235; PubMed Central PMCID: PMC2715683.

32. Velagapudi R, Hsueh YP, Geunes-Boyer S, Wright JR, Heitman J. Spores as infectious propagules of *Cryptococcus neoformans*. Infect Immun. 2009;77(10):4345–55. Epub 2009/07/22. doi: 10.1128/IAI.00542-09. PubMed PMID: 19620339; PubMed Central PMCID: PMC2747963.

33. Kwon-Chung KJ, Edman JC, Wickes BL. Genetic association of mating types and virulence in *Cryptococcus neoformans*. Infect Immun. 1992;60(2):602–5. doi: 10.1128/iai.60.2.602-605.1992. PubMed PMID: 1730495; PubMed Central PMCID: PMC257671.

34. Zhao Y, Lin J, Fan Y, Lin X. Life Cycle of *Cryptococcus neoformans*. Annu Rev Microbiol. 2019;73:17–42. Epub 20190513. doi: 10.1146/annurev-micro-020518-120210. PubMed PMID: 31082304.

35. Sun S, Coelho MA, David-Palma M, Priest SJ, Heitman J. The evolution of sexual reproduction and the mating-type locus: links to pathogenesis of *Cryptococcus* human pathogenic fungi. Annu Rev Genet. 2019;53:417–44. Epub 2019/09/21. doi: 10.1146/annurev-genet-120116-024755. PubMed PMID: 31537103; PubMed Central PMCID: PMC7025156.

36. Hull CM, Heitman J. Genetics of *Cryptococcus neoformans*. Annu Rev Genet. 2002;36:557–615. Epub 2002/11/14. doi: 10.1146/annurev.genet.36.052402.152652. PubMed PMID: 12429703.

37. Idnurm A. A tetrad analysis of the basidiomycete fungus *Cryptococcus neoformans*. Genetics. 2010;185(1):153–63. Epub 20100215. doi: 10.1534/genetics.109.113027. PubMed PMID: 20157004; PubMed Central PMCID: PMC2870951.

38. Yadav V, Sun S, Heitman J. Uniparental nuclear inheritance following bisexual mating in fungi. eLife. 2021;10. Epub 2021/08/03. doi: 10.7554/eLife.66234. PubMed PMID: 34338631; PubMed Central PMCID: PMC8412948.

39. Yan Z, Xu J. Mitochondria are inherited from the *MAT*a parent in crosses of the basidiomycete fungus *Cryptococcus neoformans*. Genetics. 2003;163(4):1315–25. doi: 10.1093/genetics/163.4.1315. PubMed PMID: 12702677; PubMed Central PMCID: PMC1462512.

40. Lavanchy G, Schwander T. Hybridogenesis. Curr Biol. 2019;29(3):539. doi: 10.1016/j.cub.2019.01.020. PubMed PMID: 30721675.

41. Casselton LA, Olesnicky NS. Molecular genetics of mating recognition in basidiomycete fungi. Microbiol Mol Biol Rev. 1998;62(1):55–70. doi: 10.1128/MMBR.62.1.55-70.1998. PubMed PMID: 9529887; PubMed Central PMCID: PMC98906.

42. Kozubowski L, Heitman J. Septins enforce morphogenetic events during sexual reproduction and contribute to virulence of *Cryptococcus neoformans*. Mol Microbiol. 2010;75(3):658–75. Epub 2009/12/01. doi: 10.1111/j.1365-2958.2009.06983.x. PubMed PMID: 19943902; PubMed Central PMCID: PMC3699866.

43. Lin X, Hull CM, Heitman J. Sexual reproduction between partners of the same mating type in *Cryptococcus neoformans*. Nature. 2005;434(7036):1017-21. Epub 2005/04/23. doi: 10.1038/nature03448. PubMed PMID: 15846346.

44. Xu J, Ali RY, Gregory DA, Amick D, Lambert SE, Yoell HJ, et al. Uniparental mitochondrial transmission in sexual crosses in *Cryptococcus neoformans*. Curr Microbiol. 2000;40(4):269–73. doi: 10.1007/s002849910053. PubMed PMID: 10688697.

45. Matha AR, Lin X. Current perspectives on uniparental mitochondrial inheritance in *Cryptococcus neoformans*. Pathogens. 2020;9(9). Epub 20200910. doi: 10.3390/pathogens9090743. PubMed PMID: 32927641; PubMed Central PMCID: PMC7559238.

46. Nielsen K, Cox GM, Wang P, Toffaletti DL, Perfect JR, Heitman J. Sexual cycle of *Cryptococcus neoformans* var. *grubii* and virulence of congenic a and alpha isolates. Infect Immun. 2003;71(9):4831–41. Epub 2003/08/23. doi: 10.1128/IAI.71.9.4831-4841.2003. PubMed PMID: 12933823; PubMed Central PMCID: PMC187335.

47. Janbon G, Ormerod KL, Paulet D, Byrnes EJ, 3rd, Yadav V, Chatterjee G, et al. Analysis of the genome and transcriptome of Cryptococcus neoformans var. grubii reveals complex RNA expression and microevolution leading to virulence attenuation. PLoS Genet. 2014;10(4):e1004261. Epub 2014/04/20. doi: 10.1371/journal.pgen.1004261. PubMed PMID: 24743168; PubMed Central PMCID: PMC3990503.

48. Arras SDM, Ormerod KL, Erpf PE, Espinosa MI, Carpenter AC, Blundell RD, et al. Convergent microevolution of *Cryptococcus neoformans* hypervirulence in the laboratory and the clinic. Sci Rep. 2017;7(1):17918. Epub 20171220. doi: 10.1038/s41598-017-18106-2. PubMed PMID: 29263343; PubMed Central PMCID: PMC5738413.

49. Arras SD, Chitty JL, Blake KL, Schulz BL, Fraser JA. A genomic safe haven for mutant complementation in *Cryptococcus neoformans*. PLoS One. 2015;10(4):e0122916. Epub 20150409. doi: 10.1371/journal.pone.0122916. PubMed PMID: 25856300; PubMed Central PMCID: PMC4391909.

50. Abramson J, Adler J, Dunger J, Evans R, Green T, Pritzel A, et al. Accurate structure prediction of biomolecular interactions with AlphaFold 3. Nature. 2024;630(8016):493-500. Epub 20240508. doi: 10.1038/s41586-024-07487-w. PubMed PMID: 38718835; PubMed Central PMCID: PMC11168924.

51. Mirdita M, Schütze K, Moriwaki Y, Heo L, Ovchinnikov S, Steinegger M. ColabFold: Making Protein folding accessible to all. Nature Methods. 2022. doi: 10.1038/s41592-022-01488-1.

52. Yadav V, Sun S, Heitman J. On the evolution of variation in sexual reproduction through the prism of eukaryotic microbes. Proc Natl Acad Sci U S A. 2023;120(10):e2219120120. Epub 20230303. doi: 10.1073/pnas.2219120120. PubMed PMID: 36867686; PubMed Central PMCID: PMC10013875.

53. Fu C, Sun S, Billmyre RB, Roach KC, Heitman J. Unisexual versus bisexual mating in *Cryptococcus neoformans*: Consequences and biological impacts. Fungal Genet Biol. 2015;78:65–75. Epub 2014/09/01. doi: 10.1016/j.fgb.2014.08.008. PubMed PMID: 25173822; PubMed Central PMCID: PMC4344436.

54. Sakuma C, Saito Y, Umehara T, Kamimura K, Maeda N, Mosca TJ, et al. The Strip-Hippo pathway regulates synaptic terminal formation by modulating actin organization at the *Drosophila* neuromuscular synapses. Cell reports. 2016;16(9):2289–97. Epub 2016/08/23. doi: 10.1016/j.celrep.2016.07.066. PubMed PMID: 27545887; PubMed Central PMCID: PMC5023852.

55. Gupta R, Kumar G, Jain BP, Chandra S, Goswami SK. Ectopic expression of 35 kDa and knocking down of 78 kDa SG2NAs induce cytoskeletal reorganization, alter membrane sialylation, and modulate the markers of EMT. Mol Cell Biochem. 2021;476(2):633–48. Epub 2020/10/22. doi: 10.1007/s11010-020-03932-2. PubMed PMID: 33083950.

56. Wernet V, Wackerle J, Fischer R. The STRIPAK component SipC is involved in morphology and cell-fate determination in the nematode-trapping fungus *Duddingtonia flagrans*. Genetics. 2022;220(1). Epub 2021/12/02. doi: 10.1093/genetics/iyab153. PubMed PMID: 34849851; PubMed Central PMCID: PMC8733638.

57. Jacinto E, Loewith R, Schmidt A, Lin S, Ruegg MA, Hall A, et al. Mammalian TOR complex 2 controls the actin cytoskeleton and is rapamycin insensitive. Nat Cell Biol. 2004;6(11):1122–8. PubMed PMID: 15467718.

58. Fan Y, Lin X. Multiple applications of a transient CRISPR-Cas9 coupled with electroporation (TRACE) system in the *Cryptococcus neoformans* species complex. Genetics. 2018;208(4):1357–72. Epub 20180214. doi: 10.1534/genetics.117.300656. PubMed PMID: 29444806; PubMed Central PMCID: PMC5887135.

59. Huang MY, Joshi MB, Boucher MJ, Lee S, Loza LC, Gaylord EA, et al. Short homology-directed repair using optimized Cas9 in the pathogen *Cryptococcus neoformans* enables rapid gene deletion and tagging. Genetics. 2022;220(1). doi: 10.1093/genetics/iyab180. PubMed PMID: 34791226; PubMed Central PMCID: PMC8733451.

60. Lin X, Chacko N, Wang L, Pavuluri Y. Generation of stable mutants and targeted gene deletion strains in *Cryptococcus neoformans* through electroporation. Med Mycol. 2015;53(3):225–34. Epub 20141224. doi: 10.1093/mmy/myu083. PubMed PMID: 25541555; PubMed Central PMCID: PMC4574871.

61. Davidson RC, Cruz MC, Sia RA, Allen B, Alspaugh JA, Heitman J. Gene disruption by biolistic transformation in serotype D strains of *Cryptococcus neoformans*. Fungal Genet Biol. 2000;29(1):38–48. Epub 2000/04/26. doi: 10.1006/fgbi.1999.1180. PubMed PMID: 10779398.

62. Tanaka R, Taguchi H, Takeo K, Miyaji M, Nishimura K. Determination of ploidy in *Cryptococcus neoformans* by flow cytometry. J Med Vet Mycol. 1996;34(5):299–301. PubMed PMID: 8912162.

