## Supplemental figures for "STRIPAK complex defects result in pseudosexual reproduction in *Cryptococcus neoformans*"

**S1 Table. Strains used in this study.**

| **Strain name** | **Description** | **Source/Reference** |
|---|---|---|
| H99α | Wild-type *MAT*α | [1] |
| KN99**a** | Wild-type *MAT***a** | [2] |
| KN99α | Wild-type *MAT*α | [2] |
| CnLC6683 | Wild-type diploid (KN99**a**/KN99α) | [3] |
| SSH118 | KN99**a** with recombinant mito. genome | [4] |
| PP71 | CnLC6683 *PPH22/pph22Δ::NAT* | [3] |
| YSB9100 | H99α *far8Δ::NAT* | [3] |
| YSB11129 | YL99**a** *far8Δ::NEO* | [3] |
| YSB11132 | YL99**a** *far8Δ::NEO* | [3] |
| PP53 | *MAT*α *pph22Δ::NAT* | [3] |
| PP55 | *MAT*α *pph22Δ::NAT* | [3] |
| PP56 | *MAT***a** *pph22Δ::NAT* | [3] |
| PP57 | *MAT***a** *pph22Δ::NAT* | [3] |
| PP58 | *MAT***a** *pph22Δ::NAT* | [3] |
| PP80 | *MAT*α *pph22Δ suppressor* (from PP53) | [3] |
| PP82 | *MAT*α *pph22Δ suppressor* (from PP55) | [3] |
| PP83 | *MAT*α *pph22Δ-8 suppressor 1* | [3] |
| PP84 | *MAT*α *pph22Δ-9 suppressor 1* | [3] |
| MCD16 | H99α *lac1Δ::URA5* | [5] |
| PP130 | PP71 *TEF1-PPG1-NEO-4* | This study |
| PP131 | PP71 *TEF1-PPG1-NEO-8* | This study |
| YSB5772 | H99α *ppg1Δ::NAT* | [6] |
| YSB5940 | H99α *ppg1Δ::NAT* | [6] |
| PP132 | *MAT*α *pph22Δ::NAT-2 TEF1-PPG1-NEO* | This study |
| PP133 | *MAT*α *pph22Δ::NAT-13 TEF1-PPG1-NEO* | This study |
| PP134 | *MAT***a** *pph22Δ::NAT-49 TEF1-PPG1-NEO* | This study |
| PP135 | *MAT***a** *pph22Δ::NAT-50 TEF1-PPG1-NEO* | This study |
| PP136 | PP71 *NOP1-GFP-HYG-7* | This study |
| PP137 | PP71 *NOP1-GFP-HYG-15* | This study |
| PP138 | *MAT***a** *pph22Δ::NAT-5 NOP1-GFP-HYG* | This study |
| PP139 | *MAT***a** *pph22Δ::NAT-7 NOP1-GFP-HYG* | This study |
| PP140 | *MAT***a** *pph22Δ::NAT-12 NOP1-GFP-HYG* | This study |
| PP141 | *MAT*α *pph22Δ::NAT-8 NOP1-GFP-HYG* | This study |
| YSC46 | KN99**a** *NOP1-GFP-HYG* | This study |
| JOHE18842 | KN99α *NOP1-mCherry-NEO* | Lab stock |
| JOHE18853 | KN99**a** *NOP1-mCherry-NAT* | Lab stock |
| JOHE10493 | KN99**a***::NEO* | Lab stock |
| PP142 | *PPH22/pph22Δ::NAT PPG1/ppg1Δ::NEO-3* | This study |
| PP143 | *PPH22/pph22Δ::NAT PPG1/ppg1Δ::NEO-11* | This study |

**S2 Table. Primers used in this study.**

| **Primer #** | **Sequence** | **Purpose** |
|---|---|---|
| JOHE53750 | CTAACTCTACTACACCTCACGGCA | *MAT***a** genotyping (*STE20***a**) |
| JOHE52751 | CGCACTGCAAAATAGATAAGTCTG | |
| JOHE52752 | GGCTGCAATCACAGCACCTTAC | *MAT*α genotyping (*STE20*α) |
| JOHE52753 | CTTCATGACATCACTCCCCTAT | |
| JOHE52754 | TGGTGGTGGTGACCCAGTTCT | Mitochondria genotyping (*COX1)* |
| JOHE52755 | CCGAAGATCTTAGGTGCCCA | |
| M13F | GTAAAACGACGGCCAG | To amplify drug resistance cassettes |
| M13R | CAGGAAACAGCTATGAC | To amplify drug resistance cassettes |
| JOHE52463 | CTGGCGGAGGATAGAAGC | *ACT1* promoter reverse primer |
| JOHE52464 | GCGAATTCGAGACAGACATCG | *TRP1* terminator forward primer |
| JOHE56046 | AATTGGGTACCGGGCCCCCCCAAGATTGTGGCTACTAT | *TEF1-PPG1-NEO* overexpression construct (cloned into pSDMA57) |
| JOHE56047 | GAAGTTTTCTGTGGAGA | |
| JOHE56048 | CGATCTCCACAGAAAACTTCATGGCACCGTTCGACCT | |
| JOHE56049 | GCGGCCGCTCTAGAAGAGAAAGTATTCGATTTG | |
| JOHE56050 | CACCGGCAGGGTATACTGTTAAGGGCCAATGAAGCACGCTGTTTTAGAGCTAGAAATAGCAAG | Safe haven gRNA |
| JOHE50175 | ACTGGTGAGTACTCAACCAAG | Safe haven screening primers |
| JOHE50176 | GGGTATGCCACAGATGCAGAT | |
| JOHE50177 | TTGGATCCTCAATTGTCTCCT | |
| JOHE50651 | GTCTTCTCCTTGTCTACAGG | To generate *ppg1::NEO* deletion construct |
| JOHE50652 | CTGGCCGTCGTTTTACTCCATTAAGCAAAGAGGGGG | |
| JOHE50653 | GTCATAGCTGTTTCCTGATACAGTACCCTGCATATCG | |
| JOHE50654 | CATGTTCTCTTTTCGCTTCC | |
| JOHE50655 | CACCGGCAGGGTATACTGTTGAACGCATCCAGCTTATTCGGTTTTAGAGCTAGAAATAGCAAG | |
| JOHE50656 | TGTACCTCATCGGCCAAAAT | *ppg1::NEO* screening primers |
| JOHE50657 | CTTCTTCTCACCCATGACCA | |
| JOHE54049 | CCACAACACATCTATCACGCGGCCGCATGGCTTTCGGTGACAGAG | GFP-tagging of *NOP1* (cloned into YSCE5) |
| JOHE54256 | ATAGAGCCACCGCCACCTGCGGCCGCAGTGTGTCGTTGGTATATGC | |

**Figure S1. Mating and genotyping analysis of progeny in *pph22*Δ x WT crosses.** A) Cell-cell fusion assay of WT::*NAT* x WT::*NEO* and *pph22*Δ::*NAT* x WT::*NEO* crosses. Cells were cocultured on MS media for three days before harvesting and plating onto selective media. Frequencies are expressed as a percentage of the wild-type control cross. Results represent three independent experiments. B) Dissection of progeny from *pph22*Δ x WT on YPD. Each row is from a separate basidium. Germinated progeny were transferred to YPD and YPD+NAT. The *pph22*Δ and wild-type parental strains were included as controls. *pph22*Δ exhibits almost no growth on YPD due to its inherent growth defects and being outcompeted for nutrients with the surrounding wild-type strains. C) Example gels from PCR genotyping showing *pph22*Δ x WT possess only one mating type and inherited mitochondria from the *MAT**a*** parent. H99α, KN99**a**, and *pph22*Δ strains served as controls.

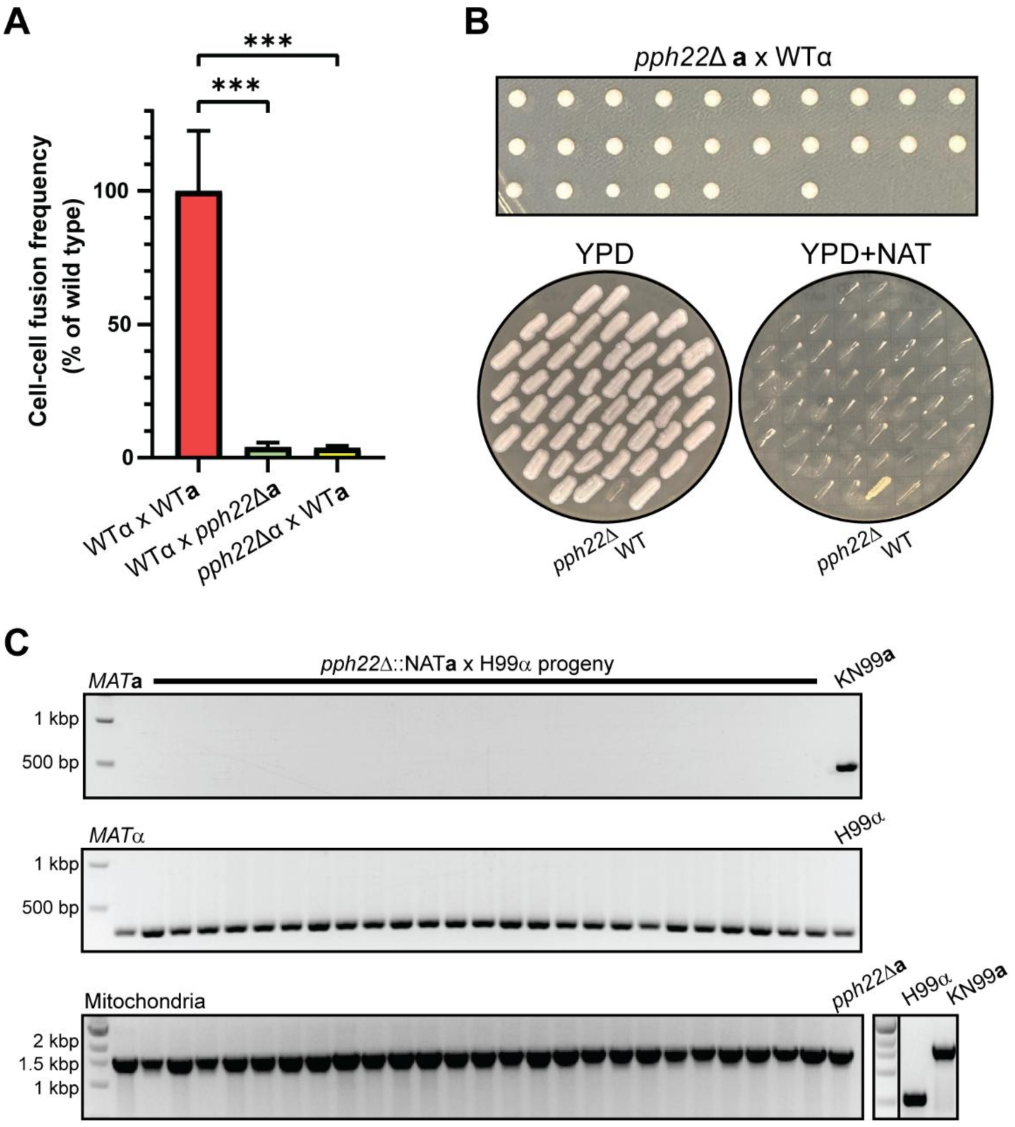

**Figure S2. Fluorescence microscopy of *C. neoformans* sexual structures.** A) Mating of wild-type and *pph22*Δ strains expressing fluorescently-tagged *NOP1* on MS media. A) WT (H99α) or *pph22*Δ expressing *NOP1-GFP* was crossed with WT (KN99**a**) expressing *NOP1-mCherry* (RFP) and mated on MS medium. B) The indicated strains expressing *NOP1-GFP* and *NOP1-RFP* were mated on MS plates for 6 to 8 weeks before imaging. DIC and fluorescence images were captured with live cells. The scale bar in each panel represents 5 µm.

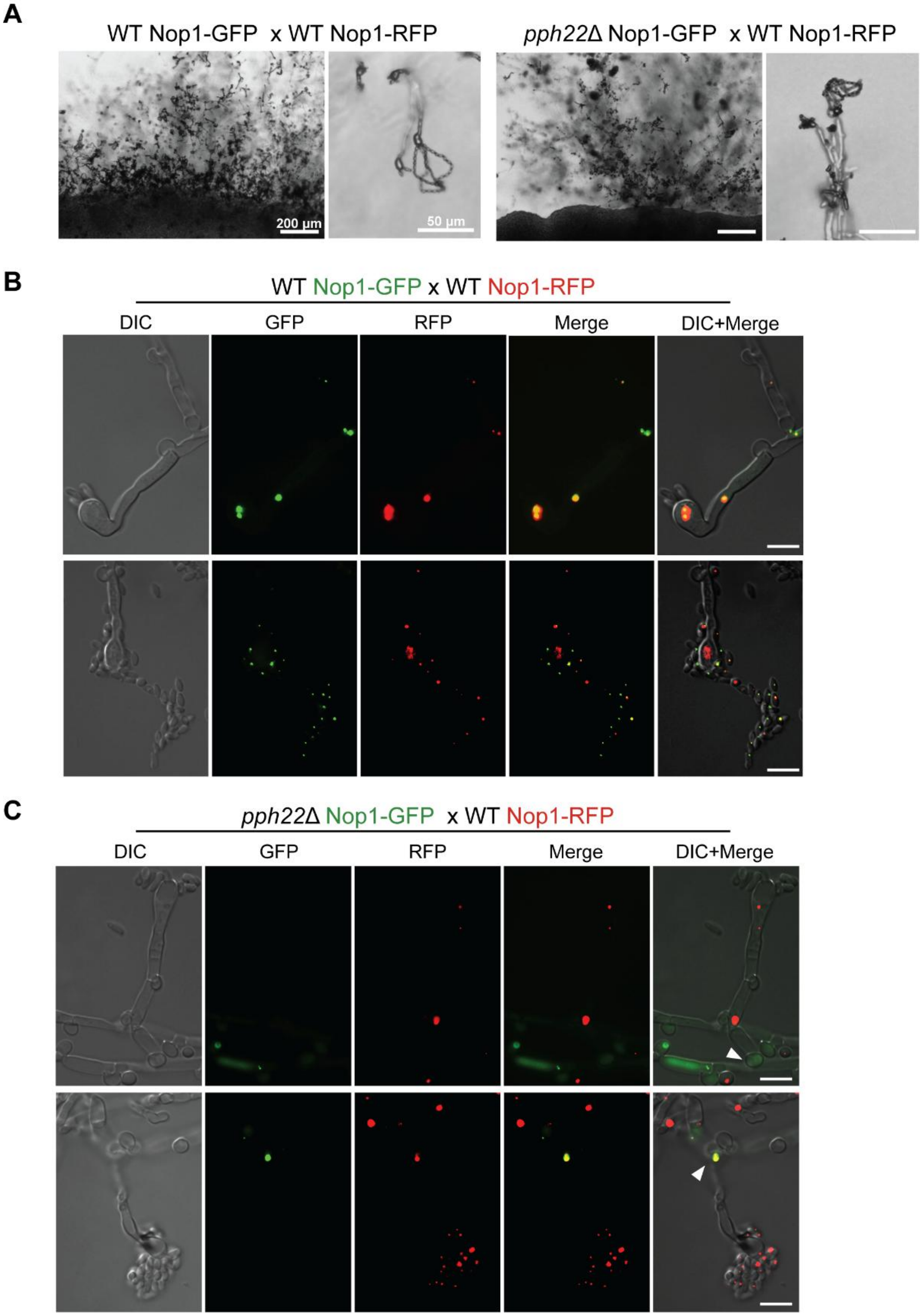

**Figure S3. Genotyping of progeny from *pph22Δ sup* x WT crosses**. A) Representative agarose gel electrophoresis analysis of PCR amplification of the *STE20* gene in *pph22Δ sup* x WT progeny shows heterozygosity of the *MAT* allele. B) Single nucleotide polymorphism in the *COX1* gene in the KN99 strain with recombinant mitochondria generates a BsrI restriction enzyme recognition site. C) Example of *COX1* fragment from PCR before and after BsrI digestion in the indicated strains used form RFLP analysis. D) RFLP following BsrI digestion of the *COX1* PCR products from *pph22Δ sup* x WT progeny. E) Genotype analysis of progeny from a third independent suppressor strain of *pph22Δ sup* crossed with wild type. F) FACS analysis of heterozygous α/**a** progeny along with KN99a (1N), CnLC6683 (2N), and PP84 as controls.

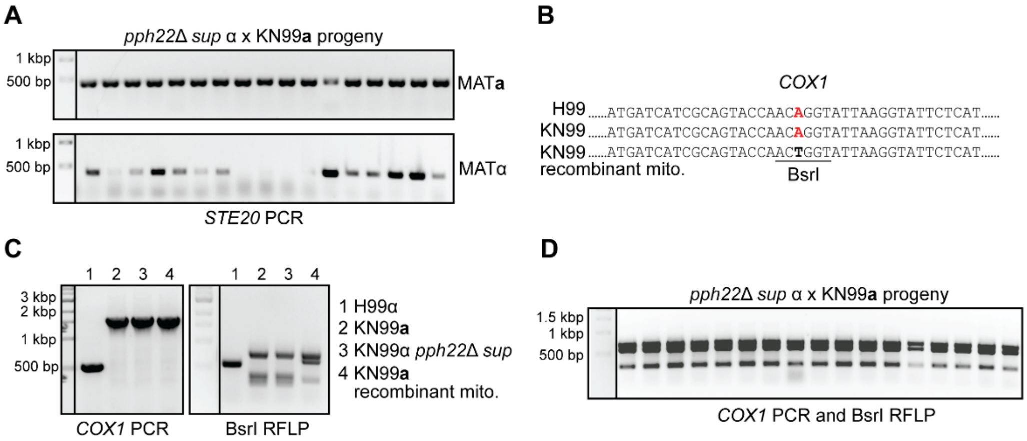


E

| Spot | Basidia # | Spores germinated/dissected | % Germination | $NAT^R$ | *MAT* | Mito. |
|---|---|---|---|---|---|---|
| *pph22Δ::NAT sup* α (PP82) x KN99**a** (SSH118) | | | | | | |
| A | 1 | 13/14 | 93 | 0 | 13 **a** | **a** |
| B | 2 | 10/13 | 77 | 0 | 10 **a** | α |
| C | 3 | 3/10 | 30 | 0 | 2 **a**, 1 α/**a** | **a** |
| D | 4 | 14/14 | 100 | 0 | 14 **a** | **a** |
| E | 5 | 13/14 | 93 | 0 | 13 **a** | **a** |
| F | 6 | 8/12 | 67 | 0 | 8 α/**a** | α |
| G | 7 | 6/10 | 60 | 0 | 3 **a**, 3 α/**a** | **a** |

F

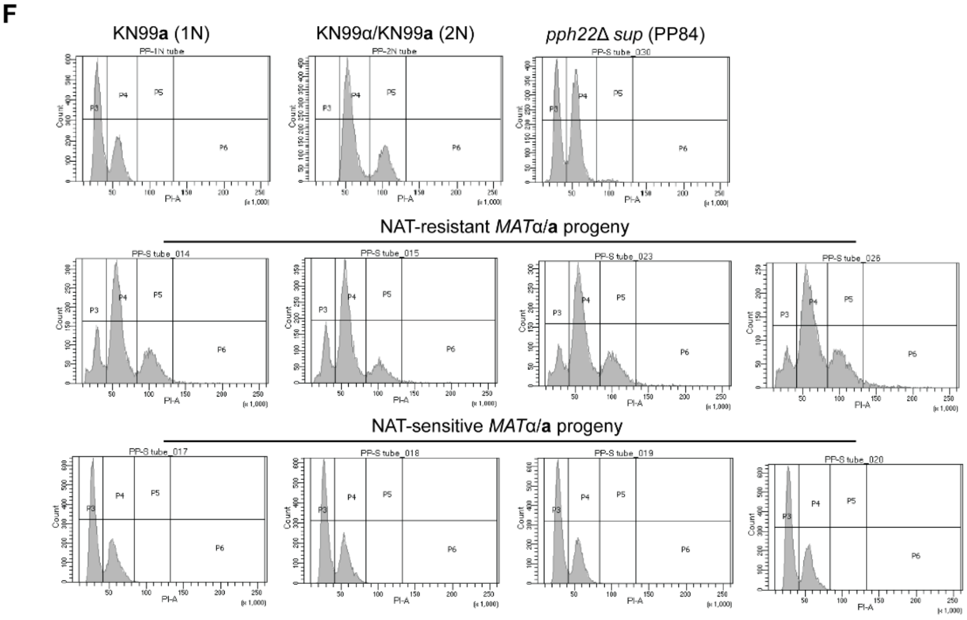

**Figure S4. AlphaFold3 multimer structure prediction of the *Cryptococcus* STRIPAK complex.** Models of STRIPAK with Pph22 (A) or Ppg1 (B) serving as the catalytic subunit. C) Homo-tetramer prediction of Far8, with protein monomers labeled A-D. The predicted multi-modular PP2A complex with Pph22 (D) or Ppg1 (E).

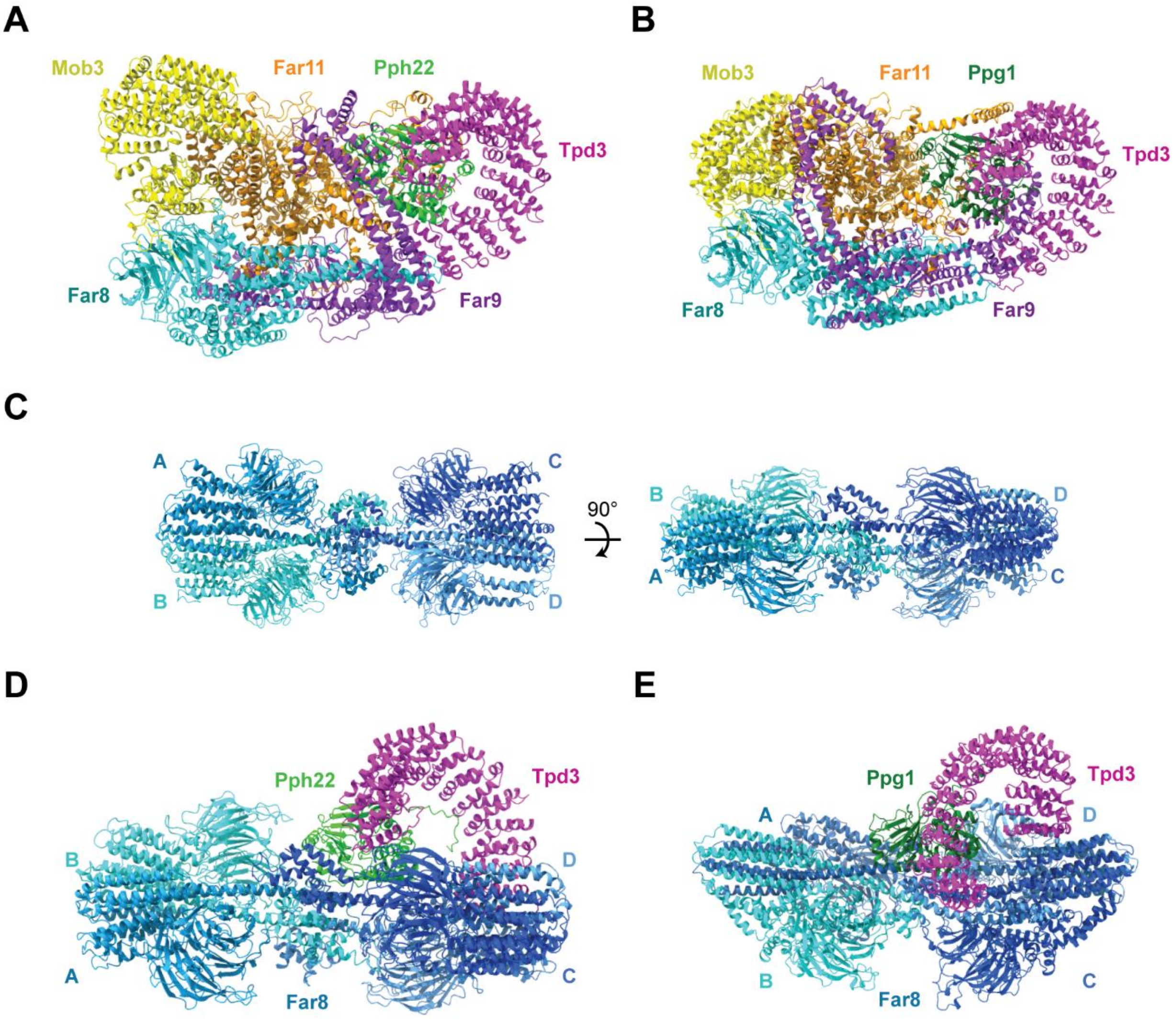

**Figure S5. Pseudosexual reproduction in STRIPAK mutants favors the wild-type parental genotype.** Distribution of genotypes among progeny in wild type, *pph22*Δ, *pph22*Δ *sup,* and *far8*Δ crosses. Number of progeny analyzed for each cross are (from left to right): 58, 282, 112, 281,122, 55. Fisher's exact test was performed for both mating type and mitochondrial type of *pph22*Δ/*pph22*Δ *sup* vs. WT and *far8*Δ vs. WT comparison groups, revealing a significant deviation in the distribution of genotypes among progeny from *pph22*Δ, *pph22*Δ *sup,* and *far8*Δ crosses compared to wild type (*P* value <0.0002, ****).

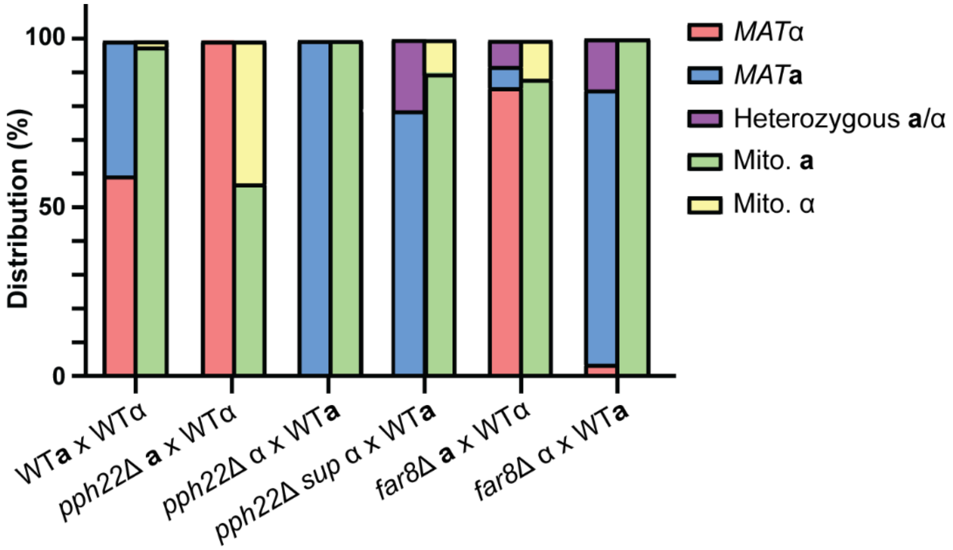

### References


1. Perfect JR, Ketabchi N, Cox GM, Ingram CW, Beiser CL. Karyotyping of *Cryptococcus neoformans* as an epidemiological tool. J Clin Microbiol. 1993;31(12):3305-9. doi: 10.1128/jcm.31.12.3305-3309.1993. PubMed PMID: 8308124; PubMed Central PMCID: PMC266409.
2. Nielsen K, Cox GM, Wang P, Toffaletti DL, Perfect JR, Heitman J. Sexual cycle of *Cryptococcus neoformans* var. *grubii* and virulence of congenic **a** and alpha isolates. Infect Immun. 2003;71(9):4831-41. Epub 2003/08/23. doi: 10.1128/IAI.71.9.4831-4841.2003. PubMed PMID: 12933823; PubMed Central PMCID: PMC187335.
3. Peterson PP, Choi JT, Fu C, Cowen LE, Sun S, Bahn YS, et al. The *Cryptococcus neoformans* STRIPAK complex controls genome stability, sexual development, and virulence. PLoS Pathog. 2024;20(11):e1012735. Epub 20241119. doi: 10.1371/journal.ppat.1012735. PubMed PMID: 39561188; PubMed Central PMCID: PMC11614259.
4. Bian Z, Xu Z, Peer A, Choi Y, Priest SJ, Akritidou K, et al. Essential genes encoded by the mating-type locus of the human fungal pathogen *Cryptococcus neoformans*. mBio. 2025:e0022325. Epub 20250225. doi: 10.1128/mbio.00223-25. PubMed PMID: 39998264.
5. Pukkila-Worley R, Gerrald QD, Kraus PR, Boily MJ, Davis MJ, Giles SS, et al. Transcriptional network of multiple capsule and melanin genes governed by the *Cryptococcus neoformans* cyclic AMP cascade. Eukaryot Cell. 2005;4(1):190-201. Epub 2005/01/12. doi: 10.1128/EC.4.1.190-201.2005. PubMed PMID: 15643074; PubMed Central PMCID: PMC544166.
6. Jin JH, Lee KT, Hong J, Lee D, Jang EH, Kim JY, et al. Genome-wide functional analysis of phosphatases in the pathogenic fungus *Cryptococcus neoformans*. Nature Communications. 2020;11(1):4212. Epub 2020/08/26. doi: 10.1038/s41467-020-18028-0. PubMed PMID: 32839469; PubMed Central PMCID: PMC7445287.
